# Steric Repulsion Counteracts ER-to-Lipid Droplet Protein Movement

**DOI:** 10.1101/2024.11.14.623598

**Authors:** Alicia Damm, Ozren Stojanović, Bianca M. Esch, Maxime Carpentier, Mohyeddine Omrane, Lionel Forêt, Florian Fröhlich, Robin Klemm, Abdou Rachid Thiam

## Abstract

Lipid droplets (LDs) are uniquely shaped organelles consisting of a neutral lipid core surrounded by a phospholipid monolayer, continuous with the cytosolic leaflet of the endoplasmic reticulum (ER). The dynamics and function of LDs are closely tied to their proteome composition, which is subject to dynamic remodeling. Key proteins essential for LD biology relocate from the ER to LDs, yet the mechanisms governing their movement and accumulation in LDs remain poorly understood. Here, we developed an innovative ex vivo tool to quantify and classify ER proteins based on their affinity for LDs. We found a broad spectrum of ER-to-LD partitioning affinities. We identified steric hindrance as a key factor in regulating ER-to-LD protein transfer, where proteins with only slightly higher LD affinity can effectively displace those with lower affinity from the LD surface. Consistent with this model, we observed that differentiation of 3T3 pre-adipocytes into adipocytes involves extensive remodeling of ER proteins targeting LDs, with Plin1—a high-affinity LD protein—becoming predominantly recruited and excluding other ER proteins. These findings highlight lateral protein-protein exclusion as a fundamental mechanism in shaping the LD proteome, providing new insights into LD biogenesis and function.

## Introduction

Lipid droplets (LDs) are dynamic organelles central to cellular lipid metabolism, with significant roles beyond lipid storage, including functions in stress responses, immunity, infection control, and cellular development ^1–4^. LDs originate from the endoplasmic reticulum (ER), where neutral lipids accumulate within the ER’s membrane bilayer. This accumulation nucleates a neutral lipid “lens” that grows and eventually buds into the cytosol ^5,6^. The core of LDs consists of stored neutral lipids, primarily triacylglycerols (TAGs) and cholesterol esters (CEs). Unlike most organelles, which are enclosed by lipid bilayers, LDs are surrounded by a phospholipid monolayer due to their unique oil-phase core ^7^. This structure forms intracellular emulsion droplets with high interfacial tension^8^, contrasting with the low surface tension of bilayer membranes. The high surface tension makes LDs conducive to recruiting amphipathic molecules, such as proteins, at their surface ^8–10^.

Due to their unique structure, LD’s surface is targeted by peripheral proteins, which are key to most LD functions ^11^. Many of these proteins are cytosolic (class II) and localize to LDs through the CYTOLD pathway (for cytosol-to-LD) via amphipathic helices (AHs), lipid anchors, or adaptor proteins ^9,12–16^. Other proteins (class I) move to the LD from the ER using the ERTOLD pathway (for ER-to-LD), typically possessing monotopic domains that allow diffusion between the ER and LD surfaces through their physical continuity ^12,13,17–21^. This includes lipid biosynthesis enzymes like Acyl-CoA synthetase 3 (ACSL3), which generates acyl-CoA for TAG synthesis ^22^, and diacylglycerol O-acyltransferase 2 (DGAT2), which converts diacylglycerol into TAG ^23^. Additionally, proteins such as Caveolin2, which are involved in caveolae formation, target newly formed LDs from the ER ^24,25^. We recently identified that monotopic proteins may also target LDs from other bilayer compartments, such as late endosomes ^26^. However, despite recent progress, the mechanisms by which monotopic proteins transfer from a phospholipid bilayer to the LD monolayer remain to be better understood ^17,18,21,27–29^.

Seipin is an integral ER protein that orchestrates LD formation. It assembles into an oligomeric ring-like structure and interacts with other monotopic proteins like LDAF1 at its center ^30–33^. Together, they form the LD assembly complex, which nucleates LDs and facilitates their growth ^30–33^. As LDs form, LDAF1 relocates from the seipin complex to the monolayer of the nascent LD ^33,34^. Similarly, ACSL3 targets early-stage LDs ^22,35^, but not all monotopic proteins follow this pathway. For instance, in Drosophila S2 cells, Glycerol-3-phosphate acyltransferase 4 (GPAT4), which catalyzes the first step in TAG synthesis, is excluded from nascent LDs and targets mature ones ^36,37^. Upon seipin deletion, GPAT4 rapidly targets nascent LDs ^38^, indicating that seipin regulates the ER-to-LD protein movement. This has also recently been evidenced by a single-molecule approach showing that the model LiveDrop peptide (*dm*GPAT4 hairpin motif ^21^) bidirectionally navigates between the ER and LD via a seipin-controlled ER-LD bridge^27^.

The CYTOLD pathway has been studied more extensively than the ERTOLD pathway, which remains far less understood. However, recent advances, particularly in Drosophila cells, have revealed a few mechanisms that regulate the ERTOLD pathway^39^. An early ERTOLD pathway, dependent on seipin, governs the initial targeting of proteins from the ER to nascent LDs . In contrast, a late ERTOLD pathway operates independently of seipin ^28^ and is promoted by the hemifusion and physical contiguity between the ER and LDs ^37,41^. The functional importance of separate ERTOLD pathways, the selectivity of monotopic proteins targeted to the LD through them, and the driving forces behind the movement of these different ERTOLD cargo proteins to the LDs remain largely unclear.

Evidence suggests that proteins possess domains by which they sense the unique physical and chemical properties of LDs. In vitro studies show in the case of TAG that hydrophobic motifs favor the LD monolayer ^17^, and higher phosphatidylcholine (PC) to phosphatidylethanolamine (PE) ratios lower the energy required for protein partitioning to LDs, likely due to the greater lipid packing defects of monolayers ^42^. The type of neutral lipid within LDs also appears to affect protein recruitment in some cases ^9,43^. Computational and mutational studies reveal that specific residues, like tryptophan and positively charged amino acids, are crucial for ER-to-LD relocation, as seen in the LiveDrop motif. Conformational changes also regulate protein recruitment, exemplified by UBXD8 and PLIN1 ^18,29,44^, which utilize membrane anchoring motifs for LD association from the ER. For instance, PLIN1’s C-terminal amphipathic helix bundle likely “unzips” to bind the LD surface ^45^.

These findings underscore the importance of lipid-membrane/protein interactions in shaping the energy landscape for protein localization to the LD monolayer versus the ER bilayer. Despite these insights, a key question remains: What factors regulate the monotopic protein partitioning level between the ER and LDs and monitor the proteome composition? Addressing these questions is crucial for understanding the occurrence and kinetics of biochemical reactions occurring at the LD surface, which underpins LD biology.

The ERTOLD pathway has primarily been studied in the context of individual proteins, focusing on structural analysis, even though free energy differences must determine their relocation extent from the ER to the LD surface. A comparative analysis of protein enrichment levels and mechanisms at LDs is still lacking. To address this, we developed a novel *ex-cellulo* tool that quantifies the ER-to-LD partitioning affinity of monotopic proteins. This approach revealed that lateral protein-protein exclusion is critical in controlling protein relocation from the ER to LDs.

## Results

### Proteins target LDs from ER with a wide range of affinities

We first asked to which extent different proteins and motifs partition between ER and LDs. To do so, we randomly picked a subset of ER proteins with different binding motifs and involved in diverse LD functions. These included LDAF1, DGAT2, AGPAT3, PLIN1, CAV2, GPAT4, FAR1, HSD17B13, ACSL3 and some of their hairpin motifs hpDGAT2, hpAGPAT3, hpGPAT4 ^21^. We included DGAT1, an ER-resident multipass transmembrane protein, as a control. For comparison, we also considered the model membrane anchors HPos and HNeu ^22^, previously developed to investigate LD-targeting ^22^. In addition, we examined the behavior of proteins from different organisms that target LDs, such as ERG6 (Saccharomyces cerevisiae) and CG2254 (Drosophila melanogaster).

HeLa cells were transfected with constructs of the target proteins fused to a fluorescent marker alongside a fluorescent marker for the ER lumen, ERox. Oleic acid (OA) was then fed to the cells for another 24 h to induce large mature LDs. This strategy focused only on the mature LD proteome and not their early biogenesis stages (Fig. 1A). We visualized the proteins through fluorescence microscopy. Fig. 1B shows three examples: (i) the hairpin of AGPAT3 is not found at LDs; (ii) ACSL3 with high intensity on both the surface of LDs and the ER; and (iii) PLIN1 shows high intensity at the LDs and virtually no signal on the ER (Fig. S1A).

**Figure 1.**
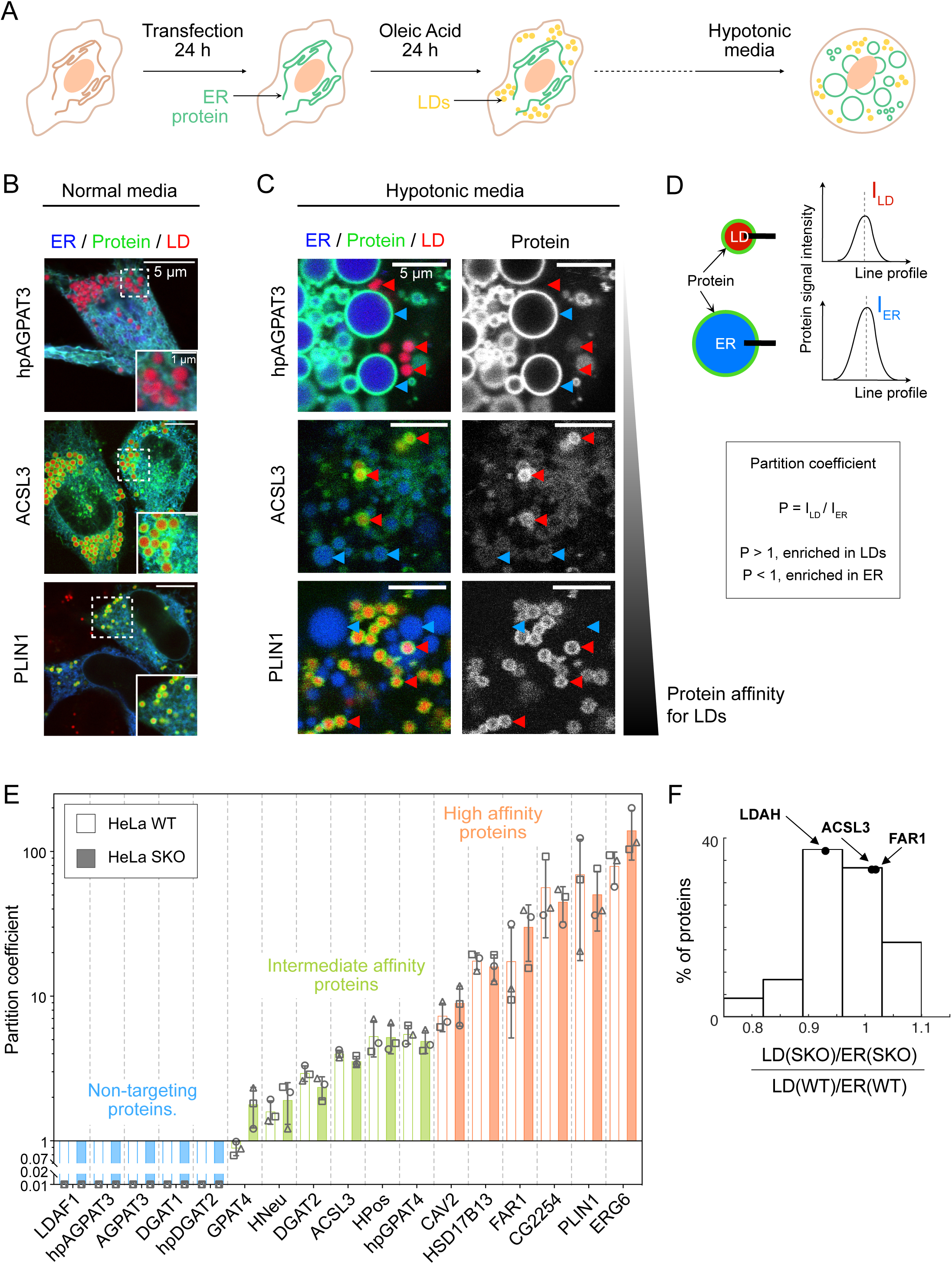
Class I proteins target LDs with a wide range of concentrations. **(A)** Schematic representation of the experimental protocol. HeLa cells are transfected with the protein of interest and an ER lumen marker for 24 h. Cells are then exposed to Oleic Acid (200 µM) for 24 h, and observed in the presence of an LD fluorescent reporter. **(B, C)** Confocal microscopy images of cells transfected with hpAGPAT3, ACSL3, PLIN1, from top to bottom. Channels: ER marker in blue (ERox), protein in green, and LD marker in red. Cells are observed in regular (B) or hypotonic media (C). In (C), red (resp. blue) arrows point to LDs (resp. ER) surface. **(D)** Quantification of the protein partition coefficient. Using a Gaussian fit, a line profile at the ER and LD equators gives respective protein intensities at the ER and LD surface. The partition coefficient is defined as the ratio between I_LD_ and I_ER_. **(E)** Partition coefficients for our 17 proteins subset are represented by mean value as a bar and standard deviation as gray whiskers, measured in HeLa WT (white bars) and HeLa SKO (colored bars). The average values of three independent experiments are represented by circle, square, and triangle gray symbols (between 3 to 5 cells for each experiment). Differences between WT and SKO partition coefficient for each protein are non-significant (unpaired t-tests; see Table 1 for values). **(F)** Distribution of the relative abundance of class I proteins in LD fraction *versus* ER-enriched fraction between HeLa WT and HeLa SKO by proteomics (27 known class I proteins were identified).

**Table 1.** P values from unpaired t test on the cells experiments (Fig. 1E) between WT and SKO measurements. T tests were performed on the average values of three independent measurements.

| Protein | P value | Summary |
| --- | --- | --- |
| GPAT4 | 0.0514 | ns |
| HNeu | 0.4617 | ns |
| DGAT2 | 0.1424 | ns |
| ACSL3 | 0.1059 | ns |
| HPos | 0.9466 | ns |
| hpGPAT4 | 0.4425 | ns |
| CAV2 | 0.4209 | ns |
| HSD17B13 | 0.5449 | ns |
| FAR1 | 0.2820 | ns |
| CG2254 | 0.5792 | ns |
| PLIN1 | 0.5922 | ns |
| ERG6 | 0.1354 | ns |

**Table 2.**
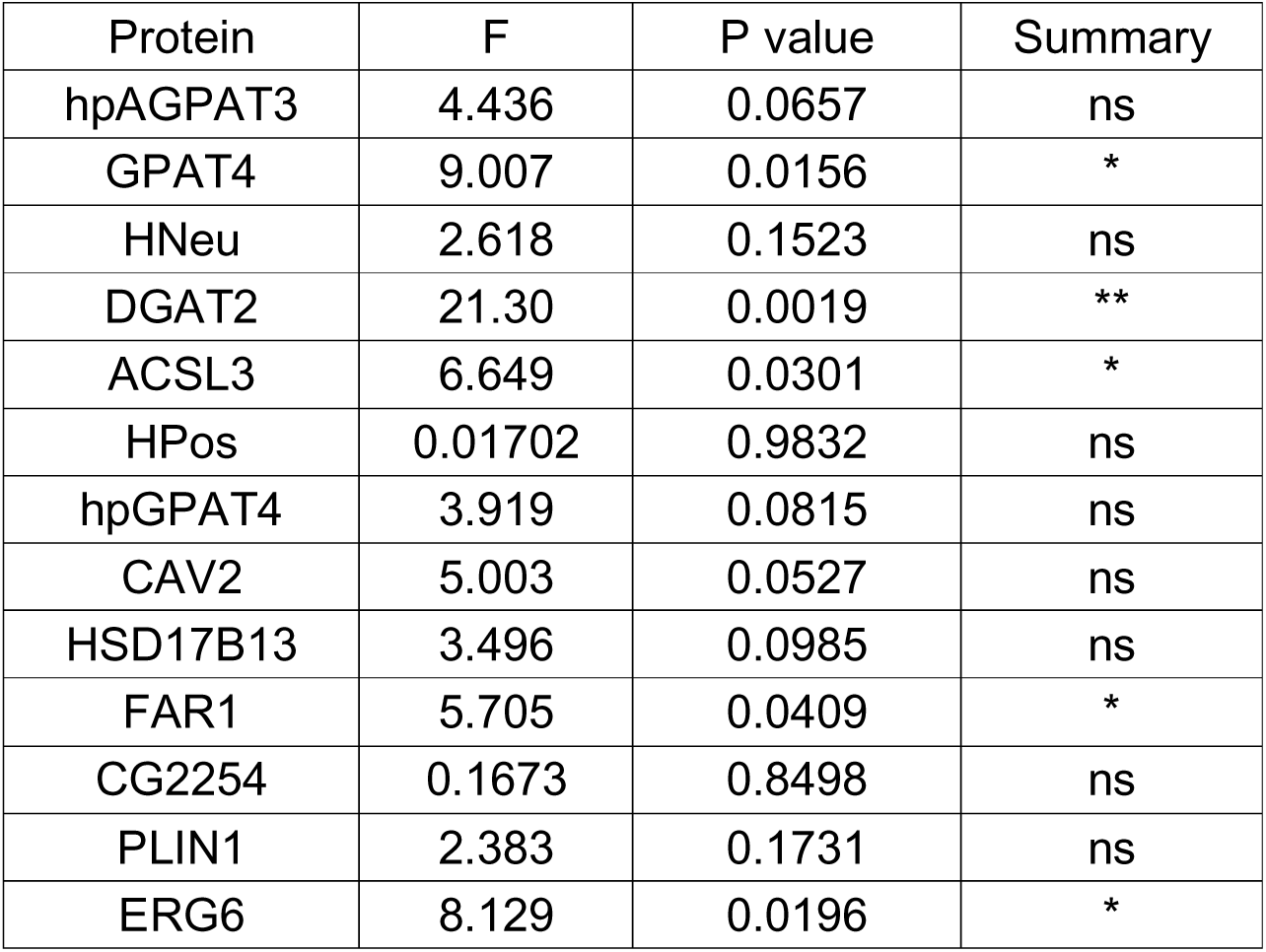
P and F values from ordinary one-way ANOVA test on the DEGERVs and cells experiments (Fig. S6B) between WT, SKO and DEGERVs measurements. One-way ANOVA were performed on the average values of three independent measurements.

| Protein | F | P value | Summary |
| --- | --- | --- | --- |
| hpAGPAT3 | 4.436 | 0.0657 | ns |
| GPAT4 | 9.007 | 0.0156 | * |
| HNeu | 2.618 | 0.1523 | ns |
| DGAT2 | 21.30 | 0.0019 | ** |
| ACSL3 | 6.649 | 0.0301 | * |
| HPos | 0.01702 | 0.9832 | ns |
| hpGPAT4 | 3.919 | 0.0815 | ns |
| CAV2 | 5.003 | 0.0527 | ns |
| HSD17B13 | 3.496 | 0.0985 | ns |
| FAR1 | 5.705 | 0.0409 | * |
| CG2254 | 0.1673 | 0.8498 | ns |
| PLIN1 | 2.383 | 0.1731 | ns |
| ERG6 | 8.129 | 0.0196 | * |

Due to the ER network structure, it is not readily possible to accurately separate the ER and LD signals. Thus, we adopted an organelle swelling approach to improve the spatial separation of different organelles, which allowed us to quantify ER vs. LD surface signals better. The osmotically-induced shape changes of most bilayer-encircled organelles ^46–49^ convert the ER into Giant ER Vesicles, GERVs ^49^ (Fig. S1B). This strategy allowed reliable and quantitative separation of the ER and LD surfaces (Fig. 1C). We determined the organelle partitioning coefficient P for each protein by calculating the ratio of intensity between the signal on the LD and the ER (Fig. 1D). When we plotted P for our 17 markers, we observed a gradual increase of the average partition coefficient across the group (Fig. 1E, S1C, and S2).

Unexpectedly, the measured partition coefficients varied over several orders of magnitude, revealing three different classes: non-targeting proteins, for which no signal was found on LDs, and we set an ultra-low partitioning coefficient of 0.01. Members in this class are LDAF1, AGPAT3, hpAGPAT3, DGAT1 and hpDGAT2. Intermediate affinity proteins such as GPAT4, HNeu, DGAT2, ACSL3, HPos, and hpGPAT4 were characterized by P<5, and high-affinity proteins with P>5 showing little to no signal on the ER were represented by CAV2, HSD17B13, FAR1, CG2254, PLIN1 and ERG6. In conclusion, this method allowed the precise assessment of the affinity of monotopic LD proteins and identified different classes within the tested ERTOLD proteins.

### Removal of seipin has a minor impact on the proteome of mature lipid droplets

Positioned at the ER-LD junction, seipin can regulate protein trafficking between these organelles, especially during the early stages of LD formation and growth^28,35,38,50^. Using our quantification method, we investigated the effect of seipin on the proteome of mature LDs by measuring the partition coefficients of the 17 selected proteins/peptides in a seipin knockout (SKO) HeLa cell line after 24 h OA loading. Interestingly, we observed no significant changes in protein affinities between wild-type (WT) and SKO cells (Fig. 1E, S2), suggesting that seipin removal does not have a significant influence on the global proteome of mature LDs at 24 h post-OA loading. This finding agrees with the hypothesis of a late ERTOLD pathway independent of seipin ^28^.

Our assessment involved only a limited number of proteins for mature LDs and, therefore, aimed to expand our analysis to the mature LD proteome. We employed mass spectrometry to assess global changes in the LD proteome between WT and SKO cells. Cells were fed with OA for a long period, after which LDs were isolated and their proteome analyzed. We quantified the abundance of monotopic ER proteins in LD fractions and normalized them to their values in ER fractions, calculating a partitioning coefficient. The distribution of the partitioning ratio between SKO and WT cells for numerous established Class I LD proteins showed a Gaussian-like distribution centered around 1, with a slight de-enrichment of approximately 20% and an enrichment of 10% (Fig. 1F). We did not find any protein that was excluded or newly recruited to mature LDs upon seipin removal. This data support also that the proteome of mature LD fractions is only marginally impacted by the absence of seipin ^51^ (Fig. S3).

We were surprised by the behavior of some of our example proteins. For example, AGPAT3, or its hairpin alone, did neither target LDs in WT nor seipin-depleted cells. DGAT2 targeted LDs equally well both in WT and SKO cells, though its hairpin alone did not target LDs in either condition. In contrast, the hairpin of GPAT4 displayed a higher affinity for LDs than the full-length protein in both WT and SKO cells, raising the possibility that there are other targeting restriction domains in the full-length protein. While most cells contained a uniform LD population, the LDs of GPAT4-transfected cells separated into two distinct populations, both in WT and SKO cells. Most LDs exhibited low levels of GPAT4 on the LD, while a subset of LDs displayed high intensity (Fig. S4), Similar observations were reported in prior studies of *Drosophila* cells ^37,38^. On average, GPAT4 showed higher relocation to LDs in SKO cells than WT cells (Fig. 1E, S4), similar to its behavior in Drosophila studies ^37,38^.

Overall, our data suggest that seipin is not a long-term regulator of the LD proteome. Only a small subset of proteins displayed slight enrichment or displacement over prolonged periods without seipin.

### Droplet-embedded giant ER vesicles mimic mature LD’s contiguity with ER

During the 24 h OA loading, one possibility is that the LD proteome undergoes significant dynamic remodeling due to factors such as protein degradation, recruitment by additional lipids and proteins, or structural changes of the ER-LD interface. As a result, the partitioning coefficient we measured may not capture the intrinsic propensity of a protein to partition between the ER and LDs. Further, although kept to a maximum of 15 min, the osmotic swelling procedure might limit the validity of our observations. To address these concerns, we developed a novel experimental system, which we term the Droplet-Embedded-GERV (DEGERV). This is inspired by previously established techniques like Droplet-Embedded Vesicles (DEV) ^6,52^ and Giant ER Vesicles (GERVs) ^49^.

DEGERVs integrate artificial lipid droplets (aLDs) within native GERVs. This approach bypasses the metabolic complexities inherent to cellular systems, allowing for the direct visualization and quantification of ERTOLD proteins, independent of factors like seipin or the structural maturation of LDs. The technique provides expansive, “empty” LD surfaces, creating a simplified environment to reveal the intrinsic targeting propensity of any ER protein toward LDs. Furthermore, DEGERVs offer insight into the determinants that guide ER proteins to LDs, free from the influence of competing metabolic processes that make identifying LD protein targeting mechanisms more difficult.

To prepare DEGERVs, HeLa WT cells were placed in a hypotonic medium for 15 minutes and lysed ^49^. The resulting GERVs were visualized by the fluorescent ERox marker (Fig. 2A) and subsequently mixed with an aLD emulsion consisting of TAG. These aLDs spontaneously fused with the GERVs ^52,53^, leading to the formation of DEGERVs (Fig. 2B).

**Figure 2.**
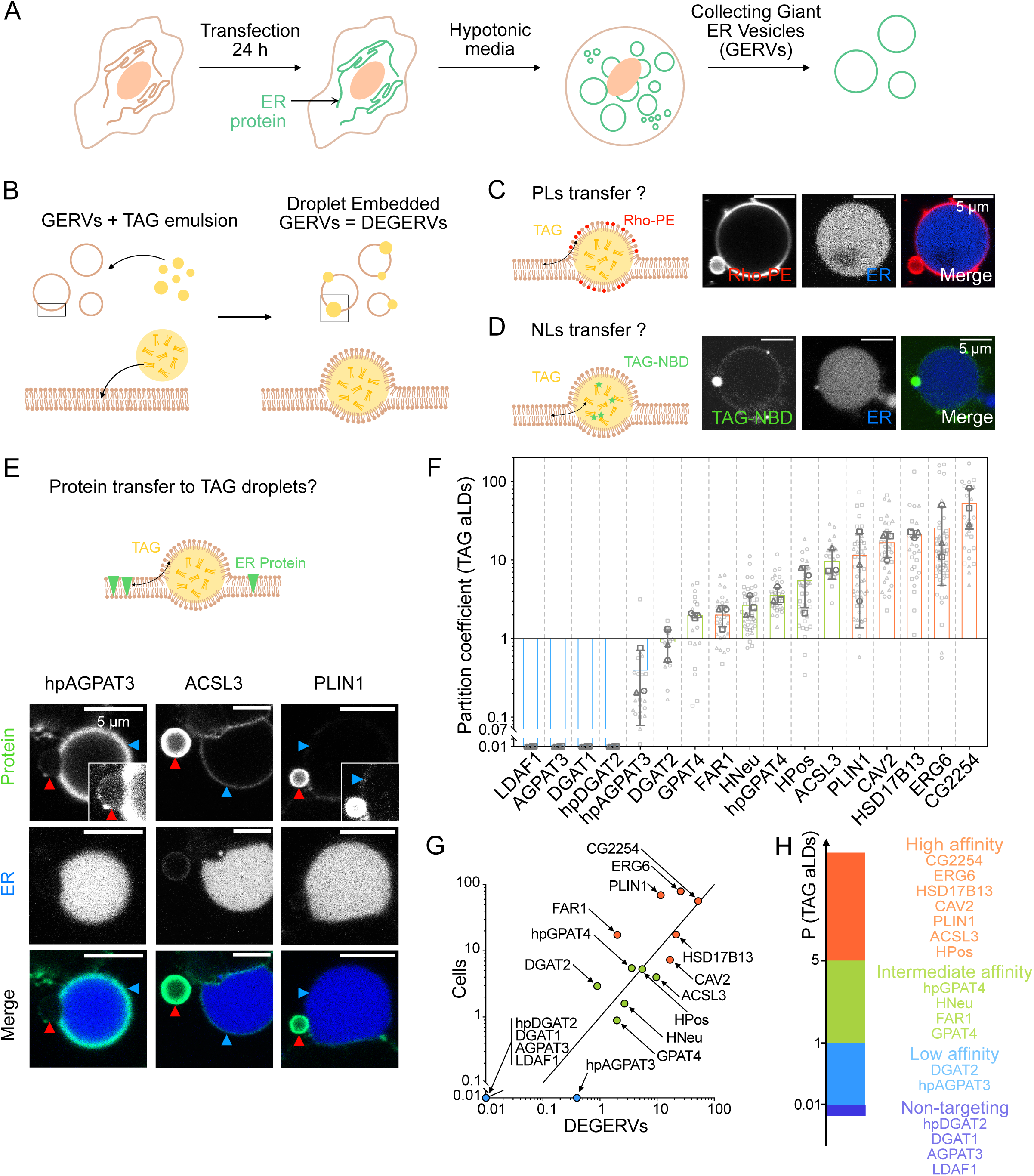
Droplet Embedded Giant ER Vesicles (DEGERVs) to dissect the regulatory mechanisms of LD targeting. **(A, B)** Schematic representation of the experimental protocol. **(A)** HeLa WT cells are transfected with the protein of interest and the ERox ER lumen marker for 24 h. Cells are then exposed to hypotonic media, and GERVs are collected as described in Material and Methods. **(B)** GERVs are mixed with a TAG-in-media droplets solution and spontaneously generate DEGERVs. **(C, D)** Characterization of TAG droplets incorporation in GERVs. **(C)** TAG-in-media emulsion was complemented with Rhodamine-DPPE (at 1:7000 w:w Rho-DPPE:TAG). Transfer of phospholipids (PLs) was observed between the TAG droplet and the GERV. **(D)** TAG-NBD-in-media emulsion is used, and a ring of NBD signal is observed at the surface of the GERV, indicating a transfer of neutral lipids (NLs) between the TAG droplet and the GERV. **(E)** Confocal microscopy images of DEGERVs prepared with cells transfected with hpAGPAT3, ACSL3, and PLIN1 (left to right). A low signal of hpAGPAT3 is observed at the TAG droplet (the inset shows a higher brightness image of hpAGPAT3 intensity on LD). In comparison, we observe a higher transfer of proteins for ACSL3 and PLIN1 (inset shows a higher brightness image of PLIN1 intensity on ER). Red arrows point to LD, and blue arrows point to ER surface. **(F)** The partition coefficient is measured for each protein, as explained in Fig. 1D. It is represented with the mean value as a bar and standard deviation as gray whiskers. The average values of three independent experiments are described as circle, square, and triangle gray symbols. All data points are represented in light gray (between 3 to 24 droplets for each experiment). **(G)** Average partition coefficient measured in cells (Fig. 1E) *versus* in DEGERVs. The plain line has a slope of 1. **(H)** Classification of proteins affinity for TAG droplets in DEGERVs.

To assess the efficient incorporation of aLDs into the GERVs, the aLDs were tagged with trace amounts of Rhodamine-DPPE (1:7000 w:w to TAG). Phospholipids from the aLDs readily transferred to the GERV membrane (Fig. 2C). FRAP experiments indicated that these phospholipids diffuse between the GERV membrane and the aLD surface, ensuring the necessary fluidity and exchange of lipids between the two compartments (Fig. S5A). Furthermore, when the aLDs were prepared with trace amounts of TAG-NBD, the fluorescent NBD signal also relocated to the GERV membrane, showing that TAG molecules were equilibrated between the aLDs and the GERV (Fig. 2D). Additional FRAP experiments revealed continuous TAG exchange between the GERV membrane and the aLDs (Fig. S5B).

Next, we employed this system to prepare GERVs from HeLa WT, which expressed our fluorescent marker proteins. DEGERVs were then formed using the same TAG-based aLDs, and protein partitioning was assessed. We observed that proteins dynamically equilibrated between the GERV membrane and aLDs, as witnessed by a fluorescence recovery of the whole aLD following photobleaching (Fig. S5C).

Most high and intermediate-affinity proteins, such as PLIN1 and ACSL3, spontaneously relocated to the aLDs (Fig. 2E). Proteins that do not specifically target LDs were generally absent from the surface of TAG-containing LDs. However, hpAGPAT3 consistently displayed a low-intensity signal on the surface of aLDs, even though this protein is not typically associated with LDs in cellular environments (inset of hpAGPAT3 image, Fig. 2E).

We determined the partition coefficients for the 17 selected proteins/peptides and plotted them according to increasing levels (Fig. 2F, Fig. S6A). As before, we set P for proteins without LD localization to 0.01. The affinity order and range were consistent between cells and DEGERV experiments, with P ranging between 0.01 and 100 (Fig. S6B). This is best illustrated by the plot of the partition coefficient in cells *vs.* DEGERVs (Fig. 2G,H), which showcases a linear correlation with a slope of 1.

There are a few notable discrepancies between protein behavior in DEGERVs and cells, revealing regulatory mechanisms that may not be functional anymore in the DEGERV system. DGAT2 and FAR1, above the correlation threshold, exhibit a lower affinity for aLDs in DEGERVs compared to their behavior in cells. This suggests that, for these proteins, a minimal TAG droplet system does not faithfully replicate the partitioning as observed in cells. It is possible that factors such as membrane curvature, lipid composition, or protein-protein interactions are different in DEGERVs than in cells, impinging on the partitioning behavior.

Conversely, proteins below the line demonstrate a higher affinity for aLDs in DEGERVs than cells. This behavior is more straightforward to interpret, as these proteins may inherently target LDs but are somehow restricted or displaced from the LD surface in cellular environments. A particularly striking example is hpAGPAT3, which is absent from LDs in cells and can yet be detected, albeit with low levels of DEGERVs. This is likely because DEGERVs initially contain no protein on the aLD surface, possibly converting some monotopic ER membrane proteins into weak ERTOLD cargoes. Once the LD surface becomes occupied, protein exclusion and competition may limit protein movement from ER to LDs.

### Competition acts as a significant regulator for protein partitioning in DEGERVs

To investigate the effect of competition between ER proteins for their relocation to LDs, we made DEGERVs from cells co-transfected with two ER proteins (Fig. 3A). We examined three distinct scenarios: (i) competition between two high-affinity proteins, ACSL3 and PLIN1, both of which exhibit partitioning coefficients (P > 5 in DEGERVs, Fig. 3B); (ii) competition between a high-affinity peptide, HPos, and an intermediate-affinity peptide, HNeu (P > 5 and P < 5, respectively, Fig. 3C); (iii) competition between a high-affinity peptide, HPos, and a low-affinity protein, DGAT2 (P < 1 in DEGERVs, Fig. 3D). In each case, we measured the partition coefficients for both proteins. We compared them to the respective P observed when they were individually expressed (Fig. 3B-D). In the presence of PLIN1, the partitioning coefficient of ACSL3 was decreased twofold (Fig. S7A), indicating that high-affinity ER proteins with different affinities can compete for targeting LD. A similar trend was seen in other high-affinity pairs, such as HPos/PLIN1 and PLIN1/HSD17B13 (Fig. S7B, S7C), further supporting this observation and indicating that PLIN1 dominates LD relocation. However, when HPos was co-transfected with intermediate-affinity HNeu (Fig. 3C) or low-affinity DGAT2 (Fig. 3D), HPos efficiently displaced HNeu or DGAT2 from the aLDs. This result demonstrates that intermediate- and low-affinity proteins are displaced from the LD surface in the DEGERV system by proteins with higher LD affinity.

**Figure 3.**
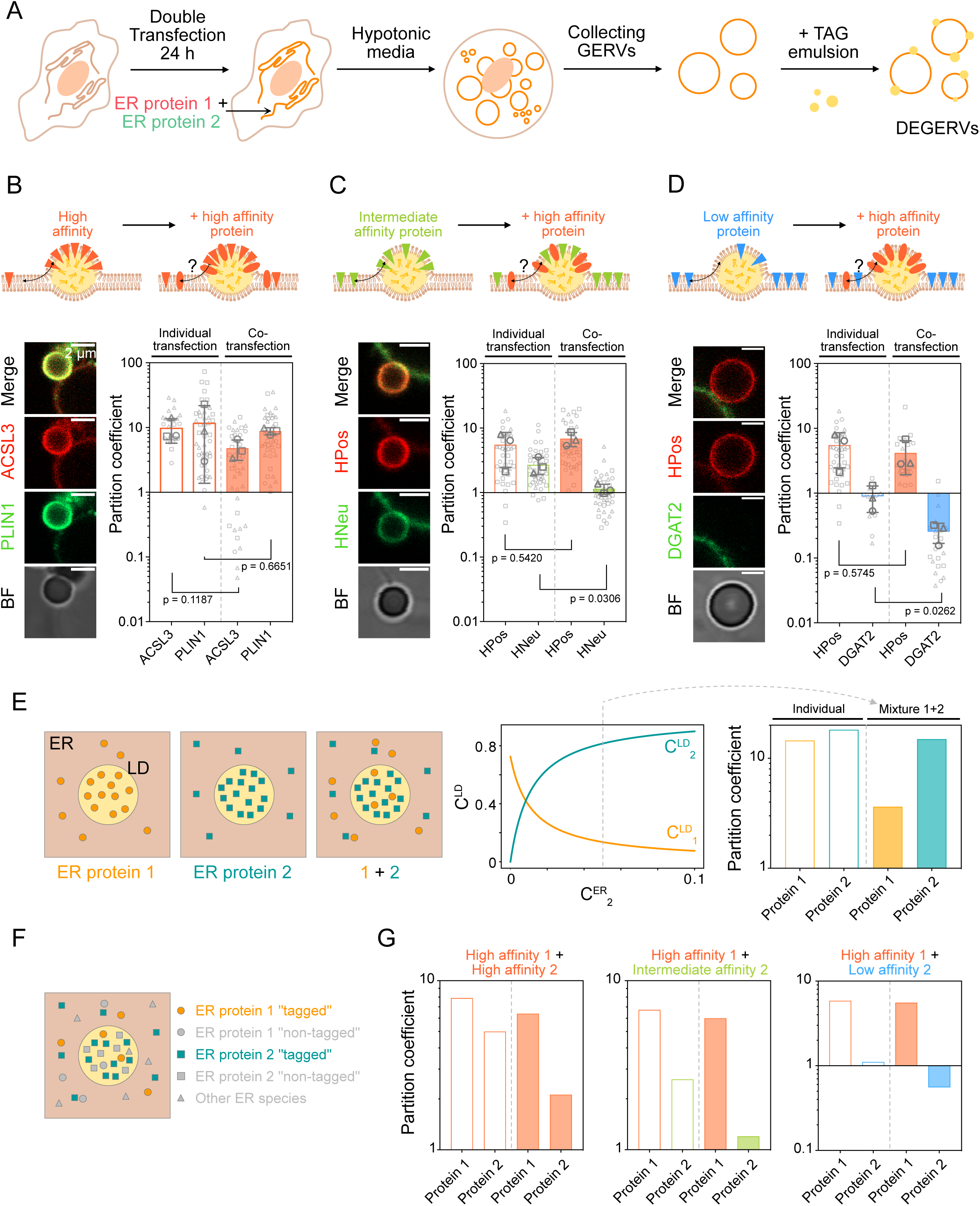
Protein competition affects LD proteome. **(A)** Schematic representation of the experimental protocol. HeLa cells are co-transfected with two proteins of interest for 24 h, then placed in hypotonic media to prepare DEGERVs. **(B, C, D)** We analyzed the competition between proteins of different affinity for LDs by measuring the partition coefficient of both proteins upon co-transfection. For each couple of proteins, we show confocal images of an embedded droplet (left panel), and we compare the partition coefficient of both proteins when transfected individually (right panel; white bars, Fig. 2F) and upon co-transfection (right panel; colored bars). We performed this experiment for a high *versus* high-affinity couple ACSL3/PLIN1 (P>5 in DEGERVs) (B), high *versus* intermediate affinity HPos/HNeu (P>5 and P<5) (C), and high *versus* low-affinity HPos/DGAT2 (P>5 and P<1) (D). The average values of three independent experiments are represented as circle, square, and triangle gray symbols. All data points are presented in light gray (between 3 to 22 droplets for each experiment). Results of the unpaired t-test is shown on the graph (see Tables 3 and 4). **(E)** A theoretical model describing steric protein-protein exclusion. For two proteins competing, the protein with the highest affinity for LDs will largely dominate LDs relocation. The protein concentration on LDs C^LD^ is plotted as a function of C^ER^ according to equation (1) with parameters: K_l_ - 50 K_2_ - 300, and C^ER^ - 0.05. Partition coefficients plotted in the right panel correspond to C^ER^ - 0.05 **(F)** To better describe our experiments and simulate the transfection of a protein, we considered a pool of all endogenous proteins present in the ER including the protein 1 (“non-tagged”), the protein 2 (“non-tagged”) and all other ER species, that will count as a protein 3 in equation (1). Transfection refers to introducing a “tagged” population of proteins 1 and 2. **(G)** Partition coefficient predicted by our model for different affinity conditions for 2 proteins expressed individually (white bars) or co-expressed (colored bars). In all conditions: C^ER^ - 0.01, C^ER^ - 0.01, C^ER^ - 0.6, K_3_ - 10, overexpression of 1 and 2 was mimicked by taking C^ER^ times 10. For high/high affinity K_l_ - 300 and K_2_ - 100 (left); for high/intermediate affinity K_l_ - 150 and K_2_ - 30 (middle); for high/low affinity K_l_ - 100 and K_2_ - 10 (right).

**Table 3.** P values from unpaired t test on the competition DEGERVs experiments (Fig. 3 and S7) between individual transfection and co-transfection measurements. T tests were performed on the average values of three independent measurements.

| Protein 1/Protein2 | P value of protein 1 | P value of protein 2 | Summary 1/2 |
| --- | --- | --- | --- |
| HPos/HNeu | 0.5420 | 0.0306 | ns / * |
| HPos/PLIN1 | 0.4260 | 0.9611 | ns / ns |
| ACSL3/PLIN1 | 0.1187 | 0.6651 | ns / ns |
| HPos/DGAT2 | 0.5745 | 0.0262 | ns / * |
| PLIN1/HSD17B13 | 0.7468 | 0.1192 | ns / ns |

**Table 4.** P values from unpaired t test on the competition DEGERVs experiments (Fig. 3 and S7) between protein 1 and 2 in individual transfection and in co-transfection measurements. T tests were performed on the average values of three independent measurements.

| Protein 1/Protein2 | P value for individual transf. | P value for co-transf. | Summary |
| --- | --- | --- | --- |
| HPos/HNeu | 0.1955 | 0.0045 | ns / ** |
| HPos/PLIN1 | 0.3792 | 0.1518 | ns / ns |
| ACSL3/PLIN1 | 0.7800 | 0.0250 | ns / * |
| HPos/DGAT2 | 0.0599 | 0.0395 | ns / * |
| PLIN1/HSD17B13 | 0.1728 | 0.4670 | ns / ns |

Therefore, the exclusion of proteins from the LD surface is primarily governed by the steric protein-protein exclusion, the thermodynamics of which is reflected in the partitioning coefficient we introduced with our system. We developed a theoretical model to assess further whether the experimental partitioning coefficient explains the displacement of proteins.

We assume proteins can freely diffuse between the LD and the ER, which we consider a protein reservoir. Each protein occupies a small area within the membrane, excluding other factors by steric repulsion. For simplicity, we assume that all proteins occupy the same space and do not interact with one another. At equilibrium, the normalized surface density of proteins *i* on the LD is (see Supplementary Materials),

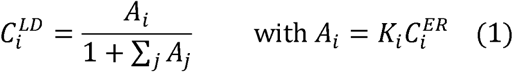

where *C^ER^_i_* is the concentration of a protein *i* in the ER and *K_i_* is the affinity constant of the LD binding reaction. The propensity of a protein to relocate to the LD surface depends on *A_i_*, the product of these two quantities. *K_i_* exponentially depends on the free energy difference of the protein between the ER and LD. If *K_i_* >1 (resp. 0 5: *K_i_* < 1), the protein preferentially relocates to the LD (resp. stays in the ER). The sum in the denominator of equation (1) accounts for the steric repulsions between proteins on LDs: it represents the combined influence of all proteins competing for space on the LD surface. The density of each protein type depends on the relative values of the parameters {*A_j_*}. To illustrate the effect of the mutual steric exclusion between two proteins, we first represented a simplified situation in which only one or two ER proteins can move onto the LDs (Fig. 3E). The simulation showed that marginal differences in the LD-affinities were sufficient to compete out the protein with lower affinity, even when both competitors had high LD-affinity by themselves.

Next, we employed this model to simulate our experiments and asked how the simultaneous incorporation of multiple proteins that relocate from the ER to LDs contributes to steric exclusion (Fig. 3F; Supplementary Material). To simulate the expression levels of the proteins, we introduced a scaling factor reflecting the protein density or concentration in the ER. Fig. 3G shows the theoretically predicted partition coefficient for different affinities of 2 proteins, expressed individually or together. Notably, the model successfully recapitulated our experimental findings using DEGERVs. In particular, proteins like DGAT2, which have a relatively lower affinity for TAG aLDs, were almost entirely excluded under certain conditions (Fig. 3D, 3G). This result underscores that competition for LD surface area through steric exclusion according to the proteins’ partition coefficient strongly influences the composition of the LD proteome. ER proteins with lower affinity for TAG aLDs, such as AGPAT3 or DGAT2, may be entirely excluded from LDs by the sole action of steric exclusion.

### The ERTOLD pathway is highly modulated to protein exclusion mechanisms

Next, we turned to DGAT2 as an example of an intermediate-affinity LD protein and investigated its behavior in more detail. We co-transfected HeLa WT cells for 24 h with DGAT2 and other ER proteins with higher LD affinity. Cells were then exposed to OA for 24 h to induce mature LDs (Fig. 4A). DGAT2 alone was enriched at LDs (Fig. 4B). When co-transfected with the HPos, DGAT2 was gradually displaced from LDs as HPos levels increased (Fig. S8A, S8B) but was never entirely excluded by HPos. Then, we tested the high-affinity protein PLIN1 against DAGT2. The overexpression of PLIN1 displaced DGAT2 from LDs back to ER almost entirely (Fig. 4C, 4D, S8C, S8D), agreeing with both our theoretical predictions and DEGERV experiments.

**Figure 4.**
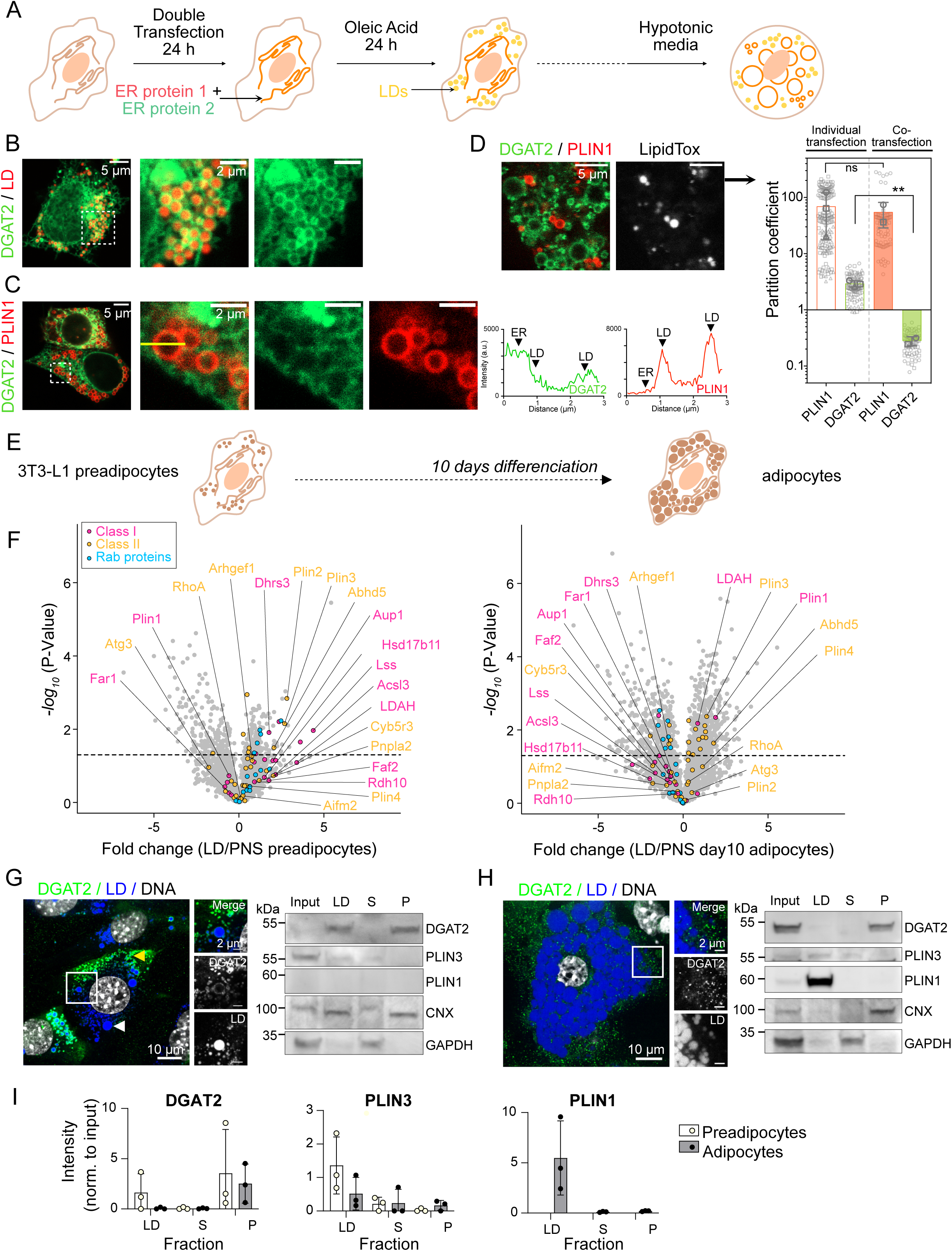
The ERTOLD pathway is highly sensitive to protein exclusion mechanisms. **(A)** Schematic representation of the experimental protocol. HeLa WT cells are co-transfected with two proteins of interest for 24 h, then exposed to Oleic Acid (200 µM) for 24 h, and observed in regular or hypotonic media. **(B)** Confocal microscopy images of HeLa WT cells transfected with DGAT2 only and using an LD marker. **(C, D)** Confocal microscopy images of cells co-transfected with PLIN1 and DGAT2 were observed in regular (C) and hypotonic (D) media. Line profiles in green and red channels along the yellow line are shown in (C). Quantification of the partition coefficients of each protein is shown in the right panel (D), comparing the partition coefficient of both proteins when transfected individually (white bars, Fig. 1E) and upon co-transfection (colored bars). Average values of three (resp. two for the co-transfection) independent experiments are represented as circle, square and triangle gray symbols. All data points are presented in light gray (between 73 to 80 droplets total, on 4 to 5 cells for each experiment). Results of the unpaired t-test are shown on the graph (see Table 5 for values). **(E)** Schematic representation of the experimental protocol using 3T3-L1 cells. **(F)** Relative abundance of proteins in LD fraction *versus* PNS fraction analyzed by proteomics in preadipocytes (left) and 10 days adipocytes (right) as a volcano plot: statistical significance -log_10_(P-Value) as a function of the fold change of the relative LD abundance of class I (pink), class II (orange) and Rab family (blue) proteins. Proteins above the dashed line correspond to a P-Value < 0.05. **(G, H)** Immunofluorescence images of endogenously expressed DGAT2 (green) in preadipocytes exposed to 500 µM oleate for 8h (G) and 10 days adipocytes (H). LDs are labelled with LipidToxRed (blue). In preadipocytes (G), DGAT2 was found to target better smaller LDs (yellow arrow) than larger ones (white arrow). Western Blots of preadipocytes (G) and 10 days adipocytes (H) after LD floatation on a sucrose cushion at 100k x g. Input is cell lysate; LD is the Lipid Droplets fraction; S is the soluble fraction, and P is the membrane pellet of a 100k x g centrifugation. GAPDH is glyceraldehyde 3-phosphate dehydrogenase and CNX, calnexin. **(I)** Quantification of WB (G, H), band intensity of DGAT2, PLIN3, and PLIN1 is measured in three independent experiments.

**Table 5.**
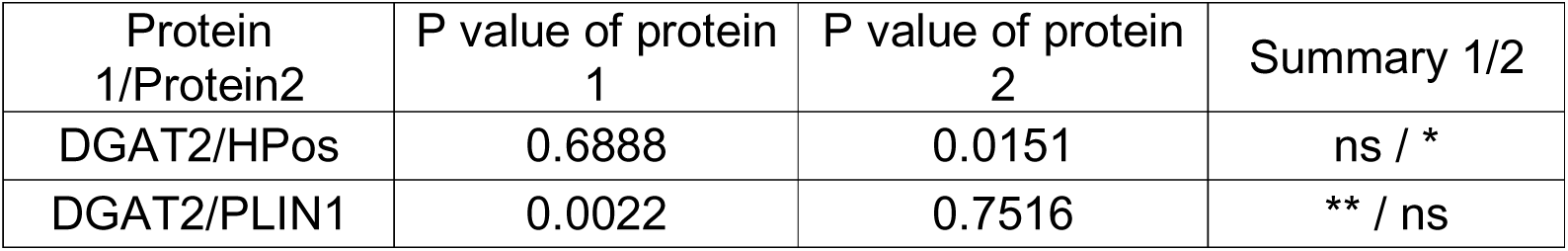
P values from unpaired t test on the competition experiments in cells (Fig. 4 and S8) between individual transfection and co-transfection measurements. T tests were performed on the average values of two or three independent measurements.

| Protein 1/Protein2 | P value of protein 1 | P value of protein 2 | Summary 1/2 |
| --- | --- | --- | --- |
| DGAT2/HPos | 0.6888 | 0.0151 | ns / * |
| DGAT2/PLIN1 | 0.0022 | 0.7516 | ** / ns |

Our analyses open the possibility that affinity-based competition between different ERTOLD cargoes may be used to tune the LD surface composition to meet cellular needs. Collectively, our theoretical and experimental results would predict that proteins with lower affinity for TAG-LDs, such as DGAT2, may, under certain circumstances, be dynamically excluded from LDs. Regulation of LD surface composition may, for instance, be achieved by controlling the expression levels of competing LD proteins. For example, PLIN1 is not expressed in pre-adipocytes, but its mRNA and protein levels increase rapidly when pre-adipocytes differentiate into adipocytes. Previous work has shown that PLIN1 efficiently excludes CYTOLD proteins such as PLIN2, PLIN3, and PLIN4 from LDs^29,45,54^, which have lower LD affinity than PLIN1. We, therefore, used adipocytes to test the validity of our model predicting the exclusion of lower affinity by high-affinity ERTOLD cargo. Specifically, we hypothesized that PLIN1 might displace lower-affinity ERTOLD cargo upon differentiation.

To compare the LD protein composition in the absence and presence of endogenously expressed PLIN1 in an unbiased fashion by proteomics, we prepared LDs from 3T3-L1 preadipocytes after fatty acid loading and from white adipocytes ten days after differentiation (Fig. 4E). The volcano plots in Fig. 4F (Fig. S9A, S9B) show significant enrichment of PLIN1 in LDs of adipocytes. Interestingly, we further confirmed our predictions, showing a global decrease in the recruitment of ERTOLD proteins after differentiation. These data may allow the interpretation that the recruitment of high-affinity PLIN1 to TAG LDs competes with other monotopic proteins.

Next, we visualized endogenously expressed DGAT2 in preadipocytes and adipocytes by immunofluorescence and determined its subcellular localization in fractionation experiments by western blotting. DGAT2 localized with high affinity to small endogenous LDs observed in untreated preadipocytes (Fig. S9C) and efficiently relocalized to LDs upon treatment with oleate (Fig. 4G). Consistently, ACSL3 was seen on both endogenous LDs as well as oleate LDs in preadipocytes. DGAT2 displayed unexpected behavior as it appeared to favor smaller LDs. ACSL3, however, accumulated all LDs, perhaps reflecting the higher position in the P-classification system. DGAT2 was virtually absent from adipocytes 10 days after differentiation. It was seen only in dot-like structures sparsely distributed over the LD surface, which may represent structures where LDs contact the ER (Fig. 4H, S9D). Subcellular fractionation validated the imaging results. In preadipocytes, DGAT2 was present in the LD fraction marked by PLIN3. Still, it was virtually absent from adipocyte LDs (Fig. 4I). A similar trend was observed for ACSL3, which has a higher affinity for LDs than DGAT2 (Fig. S9E-H). Surprisingly, ACSL3 was – just like DGAT2 – was largely excluded from adipocyte LDs. As expected, PLIN1 was not expressed in preadipocytes but was most abundant in the adipocyte LDs (Fig. 4G, 4H, 4I), likely competing with the other ERTOLD cargo.

These findings further support a mechanism by which high-affinity proteins such as PLIN1 may compete with lower-affinity ERTOLD proteins like DGAT2, possibly by steric exclusion between different ERTOLD factors.

## Discussion

Obtaining an accurate inventory of LD proteins is challenging due to the diversity in LD size, composition, and metabolic state^11^. A dynamic view of the LD proteome is essential as LDs evolve from nucleation to maturation, performing various functions across different timescales and metabolic conditions. Importantly, protein targeting to LDs is not binary; proteins accumulate at different concentrations on LD surfaces, influencing the rate of LD-related processes. Our study highlights the role of steric repulsion in controlling protein concentration and non-selective recruitment^16^ to LDs. Protein affinity for LDs depends on the energy difference between the LD monolayer and the ER bilayer, with high-affinity proteins having priority in targeting LDs—unless regulated by an energy barrier, potentially mediated by seipin during the early stage of LDs. This energy balance shifts with the cell’s metabolic state and may vary by cell type.

We classified the affinities of ER proteins targeting LDs under energy-rich conditions, with cells exposed to high concentrations of OA for 24 h. This represents a prolonged, steady-state condition compared to the shorter LD growth period. Since even slight differences in affinity can lead to significant protein exclusion (Fig. 3, S7), and the measured affinities vary by several orders of magnitude, cells must employ regulatory mechanisms to maintain LD proteome plasticity, allowing them to adapt to environmental fluctuations effectively.

Protein levels in LDs must follow a precise timeline to ensure proper metabolic responses. Early (biogenesis) and late (mature LD) ERTOLD pathways have been identified in Drosophila cells ^28^, and similar mechanisms may exist in mammalian cells. In the early pathway, proteins move from the ER to LDs by passing through the seipin complex. In the late pathway, seipin is absent at the ER-LD connection, allowing proteins like GPAT4, which fail to target nascent LDs, to use this route ^28,38^. Our findings suggest that seipin has a marginal impact on the proteome of steady-state LDs, supporting the idea of a late pathway in mammals. We speculate that seipin mainly protects the LD proteome during early growth, ensuring lipid-metabolizing enzymes, such as DGAT2, can drive LD biogenesis and maturation without being displaced by high-affinity proteins. While DGAT2 may initially target early LDs, it could later be displaced to the ER as LDs mature. High-affinity proteins, recruited later, may stabilize LDs by controlling further protein entry. This aligns with our observations of DGAT2 localization in pre-adipocytes and adipocytes. In pre-adipocytes, DGAT2 promotes LD growth by converting DAG to TAG ^36^. However, DGAT2 is excluded from the larger PLIN1 LD subpopulation in adipocytes, where growth is driven by the PLIN1/CIDE-dependent fusion ^55,56^ of smaller LDs rather than DGAT2 activity.

The regulation of protein relocation to the LD surface is critical for controlling LD function. This regulation is likely a combination of genetic coding and metabolic control, where cells activate the translation of specific proteins under particular conditions or cues. The affinities of proteins for LDs may be inherently programmed into their genetic expression profiles, enabling cells to adjust LD-associated protein levels in response to various demands, such as those encountered during the cell cycle or under stress conditions like metabolic or chronic stress. However, protein affinity for LDs is not solely determined by transcriptional control; post-translational modifications, such as phosphorylation or palmitoylation, play a crucial role in fine-tuning these interactions in response to environmental signals ^37,57–59^. For example, during nutrient deprivation, phosphorylation of PLINs may reduce their affinity for LDs, facilitating the recruitment of lipolytic enzymes necessary for lipid mobilization ^59,60^. This genetic expression and post-translation modification allow cells to regulate LD function dynamically.

Finally, when comparing our results between cells and DEGERVs, we noticed some discrepancies that warrant further investigation. For proteins like DGAT2 and FAR1, a minimal TAG LD system did not fully replicate the LD relocation observed in cells. This suggests that additional factors may influence protein affinity for LDs. Potential contributors include altered LD lipid chemistry, the highly curved bilayer-LD neck acting as an energy barrier for diffusion, and other retention factors within the ER. The DEGERV platform provides an ideal system to fine-tune these properties further and better explore their impact on the ERTOLD pathway.

## Supporting information

Supplementary informations model

## Material and Methods

### Cell Culture

HeLa WT and SKO cells (gifted by Robert Yang) were maintained in High Glucose (4.5 g/l) with stabilized Glutamine and with Sodium Pyruvate Dulbecco’s modified Eagle’s Medium (DMEM) (Dutscher) supplemented with 10% fetal bovine serum and 1% penicillin/streptomycin (GibcoBRL). Cells were cultivated 48 h at 37°C with 5% CO2. To induce feeding conditions and lipid droplet formation, cells were incubated for 24 h with DMEM supplemented with fatty acids conjugated to bovine serum albumin (BSA) (1% vol/vol), used at a concentration of 200 µM of oleic acid. Cells were regularly tested and no contamination for mycoplasma was detected.

### Cell transfections and plasmids

Cells were seeded in MatTek 3.5 mm coverslip bottom dishes (MatTek Corp. Ashland, MA) for 24 h before transfections. Cells were transfected with 2 µg of indicated plasmid using jetPEI transfection reagent (PolyPlus #10110N). Cells were transfected with different plasmids fused with fluorescent protein constructs 24 h, before 24 h feeding with OA for cells experiments (Fig. 1 and 4), or before giant organelles collection for DEGERVs experiments (no feeding, Fig. 2 and 3). Here is the list of the used plasmids.

- ERoxBFP was a gift from Erik Snapp (Addgene plasmid # 68126; http://n2t.net/addgene:68126; RRID:Addgene 68126)
- EGFP-DGAT1, EGFP-DGAT2 and mCherry-AGPAT3 were gifts from Pr. Robert Yang (University of New South Wales, Sydney, Australia)
- EGFP-FAR1 and ACSL3-mCherry were gifts from Dr. Joachim Füllekrug (Heidelberg University, Heidelberg, Germany)
- EGFP-hpAGPAT3 and EGFP-hpDGAT2 were designed and purchased from VectorBuilder (Germany)
- EGFP-PLIN1 and mCherry-PLIN1 were gifts from Pr. David B. Savage (University of Cambridge, Cambridge, United-Kingdom)
- ERG6-EGFP was a gift from Mike Henne (UT Southwestern Medical Center, Dallas, TX, USA)
- HSD17B13-turboGFP was a gift from Yaron Rotman (NIDDK, ML, USA)
- LDAF1-turboGFP was a gift from Maria Bohnert (University of Münster, Münster, Germany)
- CAV2-EGFP, EGFP-HNeu and mOrange-HPos were gifts from Albert Pol (Universitat de Barcelona, Barcelona, Spain)
- EYFP-CG2254 was a gift from Dr. Mathias Beller (Heinrich Heine University, Düsseldorf, Germany)
- mApple-hpGPAT4 was a gift from Sarah Cohen (University of North Carolina at Chapel Hill, Chapel Hill, NC, USA)
- EGFP-GPAT4 was a gift from Pr. Sander Kersten (Cornell University, Ithaca, NY, USA).

### GERVs production and extraction

After transfection (and OA feeding for cell experiments), the cultured cells were transferred into a hypotonic culture media DMEM:H2O (5:95% v/v) at pH 7.4, at 37°C, 5% CO2, for 15 min, to induce cell volume increase and GOVs. For cell experiments (Fig. 1 and 4), observation was performed at this step directly in the MatTek coverslip bottom dishes. For DEGERVs experiments (Fig. 2 and 3), cells were mechanically lysed by extensive pipetting, and the solution was filtered at 10 µm to extract GOVs, including GERVs.

### DEGERVs production

Droplets were made using an oil-in-hypotonic media emulsion: 8 μL of triolein (TO) were mixed with 300 μL of hypotonic media. The solution is subsequently sonicated and vortexed to form small droplets. The emulsion is mixed with the GERVs solution by gentle pipetting and incubated on the bench for 10 min to form DEGERVs (Fig. 2B). For the phospholipid transfer experiment (Fig. 2C), a dried film of Rhodamine-DPPE was mixed in triolein at 1:7000 w:w Rho-DPPE:TAG, prior to DEGERVs preparation. For the neutral lipid transfer experiment (Fig. 2D), a solution of TAG-NBD:TAG 1:50 v:v was used to prepare the emulsion and DEGERVs. With all approaches, the DEGERVs sample was then placed on a glass coverslip pre-treated with 10% (w/w) BSA and washed five times with hypotonic media, and it was then observed by confocal fluorescence microscopy.

### Fluorescent probes

For cells experiments, lipid droplets were tagged with HCS LipidTox™ Deep Red Neutral Lipid Stain (0.05% v/v; Cat# H34477 Thermo Fischer Scientific) for EGFP and EYFP tagged proteins, or Bodipy™ 493/503 (0.05% v/v; Thermo Fischer Scientific) for mOrange, mApple and mCherry tagged proteins.

### Confocal microscopy imaging

All micrographs were made on a Carl ZEISS LSM800, with an oil-immersed ×63 objective. EGFP, EYFP, NBD and Bodipy™ 493/503 fluorescence was excited at 488 nm, and emission was detected between 510 and 550 nm, while mOrange, mApple and mCherry tagged protein fluorescence was excited at 561 nm, and emission was detected between 580 and 650 nm. Deep red fluorescence was excited at 640 nm and was detected above 650 nm. BFP fluorescence was excited at 405 nm and detected below 500 nm. For both cells and DEGERVs experiments, Z-stack were acquired with a stack gap between 0.5 and 0.8 µm in order to cover the whole object of interest. All fluorescence signals were analyzed with ImageJ (see Data analysis section below).

### FRAP experiment in DEGERVs

Fluorescence recovery after photobleaching (FRAP) experiments were performed by bleaching either the total surface of a droplet or a part of the ER bilayer (Supplementary Fig. S5). Fluorescent proteins or lipids were bleached, and then, the recovery of signals was monitored. The FRAP curves were normalized by the fluorescence before bleaching and just after the bleach in the region of interest.

### Data analysis

All data acquired were analyzed with ImageJ. A 5 pixels-thick line is plot at the ER and LD equator lines; the line profiles obtained at these lines is fitted by a gaussian which maximum gives respectively protein intensities at ER and LD surface. For cell experiments (Fig. 1 and 4), intensities on at least 8 ER-vesicles are averaged to measure I_ER_ of one cell; and individual LD intensities I_LD_ are measured (at least 10 LDs per cell). For DEGERVs experiments (Fig. 2 and 3), I_ER_ was averaged from measurements on 5 images close to the GERV equator; and individual LD intensities I_LD_ are measured. Partition coefficient is defined as the ratio between I_LD_ and I_ER_ and can be measured for individual droplets. For proteins that didn’t target LDs at all, we manually set the partition coefficient P = 0.01. Background intensity (measured on a square of 100 pix^2^) was subtracted from I_ER_ and I_LD_.

### Statistical analysis

Statistical comparisons were made on the mean values of the three independent experiments, using a non-parametric t test or a one-way ANOVA (GraphPad Prism; unless p value is directly written on graph, ** indicates p < 0.001, * indicates p < 0.05, ns indicates p > 0.05).

### Data representation

GraphPad Prism was used to represent most graphs. Most of them are represented in log scale with mean value as a bar and standard deviation as gray whiskers. Average values of three independent experiments are represented as circle, square and triangle gray symbols; and all individual data points are represented in light gray symbols (respectively circles, squares and triangles).

### Theoretical model

See Supplementary Informations.

### Preadipocytes and adipocytes cell culture

3T3-L1 pre-adipocytes were purchased from ATCC (CL-173) and cultured at sub-confluency in DMEM with 4.5 g/L glucose and L-glutamine (Thermo Fischer Scientific, #41965062), supplanted with 10% calf serum (Sigma-Aldrich, #12133C).

For the differentiation of 3T3-L1 pre-adipocytes into adipocytes, the cells were seeded to grow to confluence the next day and then kept at confluency for 48 hours. The differentiation was induced by the addition of adipocyte differentiation medium: DMEM supplemented with 10% FBS (Sigma-Aldrich, #F7524), 172 nM (1:10000) bovine insulin (Sigma-Aldrich, #I0516), 500 µM 3-Isobutyl-1-methylxanthine (IBMX, Sigma-Aldrich #I5879), and 1 µM dexamethasone (Gbiosciences, #API-04). After 48 hours, the differentiation medium was replaced with DMEM, 10% FBS and 172 nM bovine insulin, which was replaced after two days with DMEM 10% FBS, and thereafter refreshed every two days. The cells were used for experiments at day 10 after start of differentiation.

### Lipid droplet floatation, WB and analysis

Lipid droplet floatation was done as in Majchrzak et al.^29^ Briefly, the cells were washed with and scrapped into ice-cold PBS, spun down at 600 g, 4°C for 5 minutes, and resuspended in 1 mL PBS supplemented with protease inhibitor and protease inhibitors (cOmplete™, Mini, EDTA-free Protease Inhibitor Cocktail; Merck). The cells were lysed by 20 passages through a bead homogenizer (16 µm bead, Isobiotec) and the lysate was clarified at 600 g, 4°C for 10 minutes. The post-nuclear supernatant was mixed with 60% sucrose/PBS (w/v) to a final concentration of 12% sucrose, transferred to an ultracentrifuge tube (5 x 41 mm, Beckman-Coulter, #344090) and centrifuged in an MLS-50 rotor (Beckman-Coulter) at 100,000 g, 4°C for 1h. The tubes were sealed, and flash frozen in liquid nitrogen. The top fraction was collected by cutting the frozen tube. The middle fraction was collected after thawing and the remaining pelleted fraction was washed once with PBS and then resuspended in PBS. The samples were analysed by western blot using standard techniques and antibodies against Acsl3 (Proteintech, 20710-1-AP), Dgat2 (Proteintech, 17100-1-AP), Plin1 (Cell Signaling Technology, 9349S), Gapdh (Abcam, ab181603) and calnexin (Abcam, ab140818); all at 1:1000 dilution. For immunofluorescence, the cells were grown on coverslips, fixed with 4% PFA, permeabilized with microscopy-grade 0.1% Triton X-100 for 10 min, and stained with the primary (1:200) and secondary (1:400, Alexa) antibodies in PBS/1% calf serum, and thereafter with LipidTOX Red (1:400).

### Proteomics

The samples were trypsin digested and purified for proteomics by PreOmics kit from iST, following manufacturer’s instructions. 5 μg of protein from the lipid droplet and the corresponding post-nuclear fraction was used as the input. The samples were processed as triplicates from three independent experiments.

LD and ER fractions from WT or SKO cells were normalized according to protein concentration (Fig 1F, S3). Peptides were generated following the protocol of the iST sample Preparation Kit (preOmics). Peptides were transferred to a glass vial, and 10 µL were used for reversed-phase chromatography on a Thermo Ultimate 3000 RSLCnano system, connected to a Q Exactive PLUS mass spectrometer (Thermo) via a nano-electrospray ion source. Separation was achieved using a 50 cm PepMap C18 Easy-Spray column (Thermo) with a 75 µm inner diameter, maintained at 40°C. Peptides were eluted with a linear gradient of acetonitrile (12-35% in 0.1% formic acid) over 80 minutes at a constant flow rate of 250 nL/min, followed by an increase to 60% over 20 minutes, and finally to 90% over 10 minutes. Eluted peptides were directly electrosprayed into the mass spectrometer. Mass spectra were acquired on the Q Exactive PLUS in data-dependent mode, alternating between full-scan MS and up to ten data-dependent MS/MS scans. The maximum injection time for full scans was 50 ms, with a target value of 3,000,000 at a resolution of 70,000 at m/z 200. The ten most intense multiply charged ions (z ≥ 2) from the survey scan were selected with an isolation width of 1.6 Th and fragmented using higher-energy collision dissociation (Olsen et al., 2007) with normalized collision energies of 27. Target values for MS/MS were set to 100,000, with a maximum injection time of 80 ms at a resolution of 17,500 at m/z 200. To avoid repetitive sequencing, dynamic exclusion was set to 20 seconds. The resulting MS and MS/MS spectra were analyzed using MaxQuant (version V2.4.11.0, www.maxquant.org ^61,62^). All samples were processed as triplicates from three independent experiments.

Proteins of the PNS and purified LDs from adipocytes and preadipocytes were digested using the iST Sample Preparation Kit (PreOmics). Resulting peptides were analyzed by LC-MS/MS using reversed-phase chromatography on a Thermo UltiMate 3000 RSLCnano system and sprayed into a TimsTOF HT mass spectrometer (Bruker Corporation, Bremen). For the chromatography, an Aurora Gen3 C18 column (25 cm x 75 um x 1.6 um) with CSI emitter (Ionoptics, Australia) at 40°C was used. Peptides were eluted from the column via a linear acetonitrile gradient from 10-35% in 0.1% formic acid for 44 minutes at a constant flow rate of 300 nL/min. Afterwards, the gradient was increased to 50% buffer B for 7 minutes followed by a 4-minute increase to 85% buffer B. Peptides were sprayed into the TimsTOF HT mass spectrometer through a Captive Spray Ion source at an electrospray voltage of 1.6 kV and 3 L/min DryGas. The mass spectrometer was operated in positive ion mode and a MS range from 100 to 1700 m/z using the PASEF scan mode. Ion mobility was ramped from 0.7 Vs/cm2 to 1.5 in 100 ms with an accumulation time set of 100 ms. 10 PASEF ramps per cycle resulted in a duty cycle time of 1.17 s with a dynamic exclusion time of 0.4 minutes. The Precursor ion charge state was limited from 0 to 5. The resulting MS and MS/MS spectra were analyzed using MaxQuant (V2.4.11.0, www.maxquant.org/ ^61,62^) and Perseus (V2.0.11.0, www.maxquant.org/perseus). Plots were performed with the R software package (www.r-project.org/; RRID:SCR_001905). All samples were processed as triplicates from three independent experiments.

## References

1. Bosch, M., Sweet, M.J., Parton, R.G., and Pol, A. (2021). Lipid droplets and the host– pathogen dynamic: FATal attraction? Journal of Cell Biology 220, e202104005.

2. Welte, M.A., and Gould, A.P. (2017). Lipid droplet functions beyond energy storage. Biochim Biophys Acta Mol Cell Biol Lipids 1862, 1260–1272. 10.1016/j.bbalip.2017.07.006.

3. Olzmann, J.A., and Carvalho, P. (2018). Dynamics and functions of lipid droplets. Nature Reviews Molecular Cell Biology, 1.

4. Walther, T.C., and Farese, R.V. (2012). Lipid droplets and cellular lipid metabolism. Annu. Rev. Biochem. 81, 687–714. 10.1146/annurev-biochem-061009-102430.

5. Thiam, A.R., and Forêt, L. (2016). The physics of lipid droplet nucleation, growth and budding. Biochim. Biophys. Acta 1861, 715–722. 10.1016/j.bbalip.2016.04.018.

6. Chorlay, A., Monticelli, L., Ferreira, J.V., M’barek, K.B., Ajjaji, D., Wang, S., Johnson, E., Beck, R., Omrane, M., and Beller, M. (2019). Membrane asymmetry imposes directionality on lipid droplet emergence from the ER. Developmental cell 50, 25–42.

7. Thiam, A.R., Farese Jr, R.V., and Walther, T.C. (2013). The biophysics and cell biology of lipid droplets. Nature reviews Molecular cell biology 14, 775.

8. Ben M’barek, K., Ajjaji, D., Chorlay, A., Vanni, S., Forêt, L., and Thiam, A.R. (2017). ER Membrane Phospholipids and Surface Tension Control Cellular Lipid Droplet Formation. Dev. Cell 41, 591–604.e7. 10.1016/j.devcel.2017.05.012.

9. Chorlay, A., and Thiam, A.R. (2020). Neutral lipids regulate amphipathic helix affinity for model lipid droplets. Journal of Cell Biology 219.

10. Thiam, A.R., and Dugail, I. (2019). Lipid droplet–membrane contact sites–from protein binding to function. Journal of Cell Science 132, jcs230169.

11. Bersuker, K., and Olzmann, J.A. (2017). Establishing the lipid droplet proteome: Mechanisms of lipid droplet protein targeting and degradation. Biochimica et Biophysica Acta (BBA)-Molecular and Cell Biology of Lipids 1862, 1166–1177.

12. Kory, N., Farese Jr, R.V., and Walther, T.C. (2016). Targeting fat: mechanisms of protein localization to lipid droplets. Trends in cell biology 26, 535–546.

13. Olarte, M.-J., Swanson, J.M., Walther, T.C., and Farese Jr, R.V. (2021). The CYTOLD and ERTOLD pathways for lipid droplet–protein targeting. Trends in biochemical sciences.

14. Čopič, A., Antoine-Bally, S., Gimenez-Andres, M., Garay, C.T., Antonny, B., Manni, M.M., Pagnotta, S., Guihot, J., and Jackson, C.L. (2018). A giant amphipathic helix from a perilipin that is adapted for coating lipid droplets. Nature communications 9, 1332.

15. Prévost, C., Sharp, M.E., Kory, N., Lin, Q., Voth, G.A., Farese Jr, R.V., and Walther, T.C. (2018). Mechanism and determinants of amphipathic helix-containing protein targeting to lipid droplets. Developmental cell 44, 73–86.

16. Kory, N., Thiam, A.-R., Farese Jr, R.V., and Walther, T.C. (2015). Protein crowding is a determinant of lipid droplet protein composition. Developmental cell 34, 351–363.

17. Caillon, L., Nieto, V., Gehan, P., Omrane, M., Rodriguez, N., Monticelli, L., and Thiam, A.R. (2020). Triacylglycerols sequester monotopic membrane proteins to lipid droplets. Nature Communications 11, 1–12.

18. Dhiman, R., Perera, R.S., Poojari, C.S., Wiedemann, H.T., Kappl, R., Kay, C.W., Hub, J.S., and Schrul, B. (2024). Hairpin protein partitioning from the ER to lipid droplets involves major structural rearrangements. Nature Communications 15, 4504.

19. Jacquier, N., Choudhary, V., Mari, M., Toulmay, A., Reggiori, F., and Schneiter, R. (2011). Lipid droplets are functionally connected to the endoplasmic reticulum in Saccharomyces cerevisiae. J Cell Sci 124, 2424–2437.

20. Markgraf, D.F., Klemm, R.W., Junker, M., Hannibal-Bach, H.K., Ejsing, C.S., and Rapoport, T.A. (2014). An ER protein functionally couples neutral lipid metabolism on lipid droplets to membrane lipid synthesis in the ER. Cell reports 6, 44–55.

21. Olarte, M.-J., Kim, S., Sharp, M.E., Swanson, J.M., Farese Jr, R.V., and Walther, T.C. (2020). Determinants of endoplasmic reticulum-to-lipid droplet protein targeting. Developmental Cell 54, 471–487.

22. Kassan, A., Herms, A., Fernández-Vidal, A., Bosch, M., Schieber, N.L., Reddy, B.J., Fajardo, A., Gelabert-Baldrich, M., Tebar, F., and Enrich, C. (2013). Acyl-CoA synthetase 3 promotes lipid droplet biogenesis in ER microdomains. J Cell Biol 203, 985–1001.

23. McFie, P.J., Banman, S.L., and Stone, S.J. (2018). Diacylglycerol acyltransferase-2 contains a c-terminal sequence that interacts with lipid droplets. Biochimica et Biophysica Acta (BBA)-Molecular and Cell Biology of Lipids 1863, 1068–1081.

24. Fujimoto, T., Kogo, H., Ishiguro, K., Tauchi, K., and Nomura, R. (2001). Caveolin-2 is targeted to lipid droplets, a new “membrane domain” in the cell. The Journal of cell biology 152, 1079–1086.

25. Morales-Paytuví, F., Fajardo, A., Ruiz-Mirapeix, C., Rae, J., Tebar, F., Bosch, M., Enrich, C., Collins, B.M., Parton, R.G., and Pol, A. (2023). Early proteostasis of caveolins synchronizes trafficking, degradation, and oligomerization to prevent toxic aggregation. Journal of Cell Biology 222, e202204020.

26. Carpentier, M., Omrane, M., Trager, J., Zouiouich, M., Shaaban, R., Prieur, X., Palard, M., El Khallouki, N., Giordano, F., and Harayama, T. (2024). Seipin Regulates Caveolin-1 Trafficking and Organelle Crosstalk. bioRxiv, 2024–09.

27. Mizrak, A., Kaestel-Hansen, J., Matthias, J., Harper, J.W., Hatzakis, N.S., Walther, T.C., and Farese Jr, R.V. (2024). Single-molecule analysis of protein targeting from the endoplasmic reticulum to lipid droplets. bioRxiv, 2024–08.

28. Song, J., Mizrak, A., Lee, C.-W., Cicconet, M., Lai, Z.W., Tang, W.-C., Lu, C.-H., Mohr, S.E., Farese Jr, R.V., and Walther, T.C. (2022). Identification of two pathways mediating protein targeting from ER to lipid droplets. Nature cell biology 24, 1364–1377.

29. Majchrzak, M., Stojanović, O., Ajjaji, D., M’barek, K.B., Omrane, M., Thiam, A.R., and Klemm, R.W. (2024). Perilipin membrane integration determines lipid droplet heterogeneity in differentiating adipocytes. Cell Reports 43.

30. Klug, Y.A., Deme, J.C., Corey, R.A., Renne, M.F., Stansfeld, P.J., Lea, S.M., and Carvalho, P. (2021). Mechanism of lipid droplet formation by the yeast Sei1/Ldb16 Seipin complex. Nature communications 12, 1–13.

31. Sui, X., Arlt, H., Brock, K.P., Lai, Z.W., DiMaio, F., Marks, D.S., Liao, M., Farese, R.V., and Walther, T.C. (2018). Cryo–electron microscopy structure of the lipid droplet–formation protein seipin. J Cell Biol 217, 4080–4091.

32. Su, W.-C., Lin, Y.-H., Pagac, M., and Wang, C.-W. (2019). Seipin negatively regulates sphingolipid production at the ER–LD contact site. Journal of Cell Biology 218, 3663– 3680.

33. Chung, J., Wu, X., Lambert, T.J., Lai, Z.W., Walther, T.C., and Farese Jr, R.V. (2019). LDAF1 and Seipin Form a Lipid Droplet Assembly Complex. Developmental Cell.

34. Castro, I.G., Eisenberg-Bord, M., Persiani, E., Rochford, J.J., Schuldiner, M., and Bohnert, M. (2019). Promethin Is a Conserved Seipin Partner Protein. Cells 8, 268.

35. Salo, V.T., Belevich, I., Li, S., Karhinen, L., Vihinen, H., Vigouroux, C., Magré, J., Thiele, C., Hölttä-Vuori, M., and Jokitalo, E. (2016). Seipin regulates ER–lipid droplet contacts and cargo delivery. The EMBO journal 35, 2699–2716.

36. Wilfling, F., Wang, H., Haas, J.T., Krahmer, N., Gould, T.J., Uchida, A., Cheng, J.-X., Graham, M., Christiano, R., and Fröhlich, F. (2013). Triacylglycerol synthesis enzymes mediate lipid droplet growth by relocalizing from the ER to lipid droplets. Developmental cell 24, 384– 399.

37. Wilfling, F., Thiam, A.R., Olarte, M.-J., Wang, J., Beck, R., Gould, T.J., Allgeyer, E.S., Pincet, F., Bewersdorf, J., and Farese Jr, R.V. (2014). Arf1/COPI machinery acts directly on lipid droplets and enables their connection to the ER for protein targeting. Elife 3, e01607.

38. Wang, H., Becuwe, M., Housden, B.E., Chitraju, C., Porras, A.J., Graham, M.M., Liu, X.N., Thiam, A.R., Savage, D.B., and Agarwal, A.K. (2016). Seipin is required for converting nascent to mature lipid droplets. Elife 5, e16582.

39. Dhiman, R., Caesar, S., Thiam, A.R., and Schrul, B. (2020). Mechanisms of protein targeting to lipid droplets: A unified cell biological and biophysical perspective. In Seminars in Cell & Developmental Biology (Elsevier).

40. Song, J., Mizrak, A., Lee, C.-W., Cicconet, M., Lai, Z.W., Lu, C.-H., Mohr, S.E., Farese, R.V., and Walther, T.C. (2021). Identification of two pathways mediating protein targeting from ER to lipid droplets. bioRxiv.

41. Thiam, A.R., Antonny, B., Wang, J., Delacotte, J., Wilfling, F., Walther, T.C., Beck, R., Rothman, J.E., and Pincet, F. (2013). COPI buds 60-nm lipid droplets from reconstituted water–phospholipid–triacylglyceride interfaces, suggesting a tension clamp function. Proceedings of the National Academy of Sciences 110, 13244–13249.

42. Chorlay, A., Forêt, L., and Thiam, A.R. (2021). Origin of gradients in lipid density and surface tension between connected lipid droplet and bilayer. Biophysical Journal.

43. Rogers, S., Gui, L., Kovalenko, A., Zoni, V., Carpentier, M., Ramji, K., Ben Mbarek, K., Bacle, A., Fuchs, P., and Campomanes, P. (2022). Triglyceride lipolysis triggers liquid crystalline phases in lipid droplets and alters the LD proteome. Journal of Cell Biology 221, e202205053.

44. Ajjaji, D., Ben M’barek, K., Boson, B., Omrane, M., Gassama-Diagne, A., Blaud, M., Penin, F., Diaz, E., Ducos, B., Cosset, F.-L., et al. (2021). Hepatitis C virus core protein uses triacylglycerols to fold onto the endoplasmic reticulum membrane. Traffic, 1–18. 10.1111/tra.12825.

45. Ajjaji, D., Ben M’barek, K., Mimmack, M.L., England, C., Herscovitz, H., Dong, L., Kay, R.G., Patel, S., Saudek, V., and Small, D.M. (2019). Dual binding motifs underpin the hierarchical association of perilipins1–3 with lipid droplets. Molecular biology of the cell 30, 703–716.

46. King, C., Sengupta, P., Seo, A.Y., and Lippincott-Schwartz, J. (2020). ER membranes exhibit phase behavior at sites of organelle contact. Proceedings of the National Academy of Sciences.

47. Jaiswal, A., Hoerth, C.H., Pereira, A.M.Z., and Lorenz, H. (2019). Improved spatial resolution by induced live cell and organelle swelling in hypotonic solutions. Scientific reports 9, 1–13.

48. Santinho, A., Salo, V.T., Chorlay, A., Li, S., Zhou, X., Omrane, M., Ikonen, E., and Thiam, A.R. (2020). Membrane Curvature Catalyzes Lipid Droplet Assembly. Current Biology 30, 2481–2494.e6. 10.1016/j.cub.2020.04.066.

49. Santinho, A., Carpentier, M., Lopes Sampaio, J., Omrane, M., and Thiam, A.R. (2024). Giant organelle vesicles to uncover intracellular membrane mechanics and plasticity. Nature Communications 15, 3767.

50. Grippa, A., Buxó, L., Mora, G., Funaya, C., Idrissi, F.-Z., Mancuso, F., Gomez, R., Muntanyà, J., Sabidó, E., and Carvalho, P. (2015). The seipin complex Fld1/Ldb16 stabilizes ER–lipid droplet contact sites. J Cell Biol 211, 829–844.

51. Fei, W., Zhong, L., Ta, M.T., Shui, G., Wenk, M.R., and Yang, H. (2011). The size and phospholipid composition of lipid droplets can influence their proteome. Biochemical and Biophysical Research Communications 415, 455–462. 10.1016/j.bbrc.2011.10.091.

52. Chorlay, A., Santinho, A., and Thiam, A.R. (2020). Making Droplet-Embedded Vesicles to Model Cellular Lipid Droplets. STAR Protocols, 100116.

53. Chorlay, A., and Thiam, A.R. (2018). An asymmetry in monolayer tension regulates lipid droplet budding direction. Biophysical journal 114, 631–640.

54. Wolins, N.E., Brasaemle, D.L., and Bickel, P.E. (2006). A proposed model of fat packaging by exchangeable lipid droplet proteins. FEBS letters 580, 5484–5491.

55. Sun, Z., Gong, J., Wu, H., Xu, W., Wu, L., Xu, D., Gao, J., Wu, J., Yang, H., and Yang, M. (2013). Perilipin1 promotes unilocular lipid droplet formation through the activation of Fsp27 in adipocytes. Nature communications 4, 1594.

56. Xu, L., Li, L., Wu, L., Li, P., and Chen, F. (2024). CIDE proteins and their regulatory mechanisms in lipid droplet fusion and growth. FEBS Letters 598, 1154–1169. 10.1002/1873-3468.14823.

57. Soni, K.G., Mardones, G.A., Sougrat, R., Smirnova, E., Jackson, C.L., and Bonifacino, J.S. (2009). Coatomer-dependent protein delivery to lipid droplets. Journal of cell science 122, 1834–1841.

58. Greenberg, A.S., Egan, J.J., Wek, S.A., Garty, N.B., Blanchette-Mackie, E.J., and Londos, C. (1991). Perilipin, a major hormonally regulated adipocyte-specific phosphoprotein associated with the periphery of lipid storage droplets. Journal of Biological Chemistry 266, 11341–11346.

59. Sztalryd, C., and Brasaemle, D.L. (2017). The perilipin family of lipid droplet proteins: Gatekeepers of intracellular lipolysis. Biochimica et biophysica acta (bba)-molecular and cell biology of lipids 1862, 1221–1232.

60. Miyoshi, H., Souza, S.C., Zhang, H.-H., Strissel, K.J., Christoffolete, M.A., Kovsan, J., Rudich, A., Kraemer, F.B., Bianco, A.C., and Obin, M.S. (2006). Perilipin promotes hormone-sensitive lipase-mediated adipocyte lipolysis via phosphorylation-dependent and-independent mechanisms. Journal of Biological Chemistry 281, 15837–15844.

61. Cox, J., and Mann, M. (2008). MaxQuant enables high peptide identification rates, individualized ppb-range mass accuracies and proteome-wide protein quantification. Nature biotechnology 26, 1367–1372.

62. Cox, J., and Mann, M. (2011). Quantitative, High-Resolution Proteomics for Data-Driven Systems Biology. Annu. Rev. Biochem. 80, 273–299. 10.1146/annurev-biochem-061308-093216.

