## Supplementary informations model for "Steric Repulsion Counteracts ER-to-Lipid Droplet Protein Movement"

### Supplementary Information.
### Simple thermodynamic model of protein partition between Endoplasmic Reticulum and Lipid Droplet

October 2, 2024

We present a simple thermodynamic model of membrane protein partitioning between two subsystems. Our goal is not to make quantitative comparisons and tests with the *in vivo* experimental results, leading in particular to the extraction of parameter values. The complexity of the real system, the uncertainties in the measurements, and the unknown parameters of the model do not allow such comparisons. Our goal is rather to present and illustrate, through the use of a well-known model kept as simple as possible, the physical phenomena of steric exclusion and its influence on protein distribution. The model predictions with plausible values of the parameters are compared at a qualitative level with the tendencies observed experimentally.

#### 1 Equilibrium condition

We consider a Lipid Droplet (LD) connected with the Endoplasmic Reticulum (ER). The continuity of the ER membrane leaflets with the LD interfaces allows some proteins to freely diffuse between the LD and the ER. At equilibrium, for each specy $i$ that can be exchanged between the ER and LD, its distribution is imposed by the balance of the chemical potentials of that protein in the ER and in the LD,

$$\mu_i^{ER} = \mu_i^{LD} \ . \tag{1}$$

To express these chemical potentials in terms of the LD and ER physico-chemical parameters, and their protein compositions, we used a classical model of 2D fluid mixture taking into account the steric repulsion between protein molecules. This model is presented in the next section.

#### 2 Model of membrane proteins mixture

The ER membrane and the LD surface covered with various proteins are both described using a classical model of 2D fluid mixture. In this model, the membrane of area $A$ containing $M$ protein species, is divided into $N_0$ sites of area $a_0$ (then $A = N_0 a_0$). Each site can be occupied by either 1 or 0 protein. The fact that a site cannot be occupied by more than one protein describes the effect of the steric repulsion between the proteins, also named "excluded area". The parameter $a_0$ accounts for the typical area occupied by a single protein and is taken to be equal for all protein species for simplicity. Other type of non-specific protein-protein interactions are neglected. We denote $N_i$ the number of molecules of type $i$ with $i \in \{1, 2, ..., M\}$, and $C_i = N_i a_0 / A = N_i / N_0$, the surface fraction of proteins of type $i$. The total number of proteins $N$ and total surface fraction of proteins $C$ are then,

$$N = \sum_{j=1}^{M} N_j \quad , \quad C = \sum_{j=1}^{M} C_j$$

The free energy of the protein mixture reads,

$$F = \sum_{i=1}^{M} \epsilon_i N_i - k_B T \left[ N_0 \ln N_0 - (N_0 - N) \ln(N_0 - N) - \sum_{i=1}^{M} N_i \ln N_i \right] \ , \qquad (2)$$

where $\epsilon_i$ is the free energy of a single protein $i$ and accounts for the interactions between the protein and its host membrane, $k_B$ is the Boltzmann constant and $T$ is the temperature. The second term under bracket in (2) is the entropy of mixing of the proteins, including the effect of the limited accessible area. This free energy is widely used in the context of phase separation modelling, see for exemple [1]. The version used here neglects the short range protein-protein interactions (except steric interactions). The chemical potential of a protein of type $i$ is then,

$$\mu_i = \frac{\partial F}{\partial N_i} = \epsilon_i + k_B T \ln \left( \frac{C_i}{1 - C} \right) \qquad (3)$$

Note that $\mu_i$ depends on the surface fraction of all the other species through the presence of $C = \sum_{j=1}^{M} C_j$ in the its expression.

For a given protein, its physical environment is not the same in the ER and in the LD. It follows that the parameter $\epsilon_i$ in (2) is not the same in the ER and in the LD. We denote $\epsilon_i^{ER}$ and $\epsilon_i^{LD}$, their value in the ER and in a LD respectively. The surface fractions of proteins are also different in the ER and in the LD, and are denoted $C_i^{ER}$ and $C_i^{LD}$.

#### 3 Protein composition of the LD at equilibrium

The composition of the ER is regulated and its size is large as compared to a LD. The ER can thus be considered as a reservoir with fixed chemical potentials. Using the equilibrium condition (1), and equation (3) to express the chemical potential of the LD species, the equilibrium surface fraction of proteins $i$ in the LD can be written,

$$C_i^{LD} = \frac{A_i}{1 + \sum_{j=1}^{M} A_j} \qquad (4)$$

Each ER specy is thus characterized by a single fixed parameter $A_i$ which reads,

$$A_i = \exp \left[ (\mu_i^{ER} - \epsilon_i^{LD}) / k_B T \right]$$

This parameter corresponds to the surface fraction of protein $i$ in the virtual situation where the steric interactions between proteins would be "switched off" *i.e.*, $C_i^{LD} = A_i$ in limit $a_0 \to 0$. The higher this parameter $A_i$, the higher the amount of protein $i$ in the LD. One can notice from (4) that the density ratio of two species is,

$$\frac{C_i^{LD}}{C_j^{LD}} = \frac{A_i}{A_j} \ .$$

The denominator in equation (4) arises from the mutual competition between the proteins for the access to the LD surface : the higher the $A_j$ with $j \neq i$, the lower the $C_i^{LD}$. Proteins of type $j$ with a high affinity for the LD (*i.e.* large $A_j$) drastically reduces the amount of proteins $i$ on the LD as compare to a situation where proteins $j$ would be absent. As a conclusion, the amount of a given protein in the LD depends on the LD affinity of all the species, which mutually exclude each others.

#### 4 Protein partitioning between the LD and ER

To relate the surface density of a protein in the LD to that in the ER, we specify the chemical potentials of the ER species. Starting from (3), the chemical potential of the protein $i$ in the ER can be written,

$$\mu_i^{ER} = \tilde{\epsilon}_i^{ER} + k_B T \ln C_i^{ER} \ ,$$

where, $\tilde{\epsilon}_i^{ER} = \epsilon_i^{ER} - k_B T \ln(1 - C^{ER})$ is a free energy per protein which includes the steric interactions. In the ER, the total surface fraction of proteins $C^{ER}$ is not low but the surface fraction of each specy $i$ is low, $C_i^{ER} \ll 1$. It follows that the parameter $\tilde{\epsilon}_i^{ER}$ can be taken as a constant independent of $C_i^{ER}$. We can then write,

$$A_i = K_i C_i^{ER} \ , \tag{5}$$

where the parameter,

$$K_i = \exp\left[(\tilde{\epsilon}_i^{ER} - \epsilon_i^{LD})/k_B T\right]$$

characterizes the relative affinity of a ER protein of type $i$ for the LD. If $\epsilon_i^{LD} > \tilde{\epsilon}_i^{ER}$ then $K_i > 1$, the proteins $i$ prefer to be on the LD surface rather than on the ER membrane and conversely. Note that the energies $\epsilon_i^{ER,LD}$ should be of the order of several $k_B T$. The parameter $K_i$ can thus be possibly as large as a few hundreds.

#### 5 Competition between two protein species

To draw comparisons with the experimental results of the paper, we applied equation (4) to the case where two ER proteins species are tagged and are indexed 1 and 2. Of course the ER contains many other untagged protein species. The equilibrium density of the tagged proteins in the LD reads (4),

$$C_1^{LD} = \frac{A_1}{1 + A_1 + A_2 + A_{ER}} \quad \text{and} \quad C_2^{LD} = \frac{A_2}{1 + A_1 + A_2 + A_{ER}} \quad \text{with} \quad A_{ER} = \sum_{j=3}^{M} A_j \ . \tag{6}$$

The effect of all the other untagged proteins is thus characterized by a single parameter $A_{ER}$. In other word, this pool of untagged proteins has the same effect as a single third protein type indexed 3, $A_{ER} = A_3 = K_3 C_3^{ER}$, where in the second equality $C_3^{ER}$ is the total surface fraction of untagged proteins and $K_3$ the average of value $K_j$ over these proteins.

**Case of a binary mixture**.
To understand the steric exclusion effect, it is first interesting to consider the fictive situation with only two proteins (no untagged proteins $A_{ER} = 0$) and compare the case with protein 1 alone ($A_2 = 0$) and both protein present. In the second case, the amount of protein 1 in the LD will be reduced by a factor $(1 + A_1)/(1 + A_1 + A_2)$ as compared to the first case. This reduction is significant if $A_2$ is ($i$) close or larger than 1, which implies a high affinity of protein 2 for the LD, and (ii) of the order or larger than $A_1$. See the histograms of Figure 3E of the main part, where we used $A_1 = 2.5$ and $A_2 = 15$. To illustrate the exclusion of the protein 1 by the protein 2, we also show in the figure 3E the evolution of $C_1^{LD}$ and $C_2^{LD}$ when the density of protein 2 in the ER, $C_2^{ER}$, is increased little by little, thereby increasing $A_2 = K_2 C_2^{ER}$. We used $K_2 = 300$ and $C_1^{ER} = C_2^{ER} = 0.05$.

**Effect of the pool of untagged proteins.**
The competition between two proteins discussed above is reduced in presence of the other ER proteins willing to go the LD and contributing to the parameter $A_{ER}$ in (6). According to equation (6), the larger this parameter $A_{ER}$ the lower the direct effect of $A_2$ on $C_1^{LD}$ (and of $A_1$

on $C_2^{LD}$). Revisiting the comparison of the preceding paragraph (protein 2 absent *vs* protein 2 present), the reduction factor of the density of protein 1 in the LD due to the presence of protein 2 is now $(1 + A_1 + A_{ER})/(1 + A_1 + A_2 + A_{ER})$. This factor deviates significantly from 1 only if $A_2$ is large or comparable to $A_{ER}$.

**Comparison with experiments.**
In the *in vivo* experiments, the ER membrane always contains the full set of proteins. The effect of the transfection is to increase by an (unknown) factor the density of the transfected proteins in the ER, thereby increasing its propensity to go the the LD, equation5). In the model, it can be mimicked by increasing $C_i^{ER}$ and thus $A_i$ with $i$ the label of the transfected protein, by an arbitrary factor. We did so for three pairs of tagged proteins with different couples of values $(A_1, A_2)$, see Figure 3G.

### References

[1] Joel Berry, Clifford P Brangwynne, and Mikko Haataja. Physical principles of intracellular organization via active and passive phase transitions. *Reports on Progress in Physics*, 81(4):046601, 2018.
