## Supplementary figures caption for "Steric Repulsion Counteracts ER-to-Lipid Droplet Protein Movement"

**Figure S1. Confocal images of transfected HeLa WT cells in normal or hypotonic media.** **(A)** Separated channels of confocal microscopy images of cells in normal media, transfected with, respectively from left to right, hpAGPAT3, ACSL3, PLIN1. **(B)** Image of a cell in hypotonic media, fluorescence of ER lumen marker (blue), ER protein (green) and LDs marker (red) on the left and bright field on the right. **(C)** Confocal images of HeLa WT cells in hypotonic media for each protein. All scale bars are 5 µm.

**Figure S2. Protein partition coefficient classification between HeLa WT and SKO cells.** Partition coefficients for our 17 proteins subset are represented by mean value as a bar and standard deviation as gray whiskers (log scale), measured in HeLa WT (white bars) and in HeLa SKO (colored bars). All data points (= individual droplets) of three independent experiments are represented as circle, square and triangle gray symbols (between 3 to 5 cells for each experiment, with at least 8 droplets analyzed per cell).

**Figure S3. Proteomics analysis between HeLa WT and SKO cells. (A)** Western Blots of cell lysates of HeLa WT and HeLa SKO separated by ultracentrifugation. First fraction corresponds to LDs (marked with PLIN3) and fractions six and seven were selected for ER fraction (marked by Calretinulin). **(B)** Relative abundance of proteins in LD fraction *versus* ER fraction analyzed by proteomics in HeLa WT (top) and HeLa SKO (bottom) as a volcano plot: statistical significance -log_10_(P-Value) as a function of the fold change of the relative LD abundance of class I (pink), class II (orange) and Rab family (blue) proteins. Proteins above the dashed line correspond to a P-Value < 0.05. **(C)** Heat map of the ratio of intensity of proteins in LD fraction between HeLa SKO and WT, analyzed by proteomics (80 relevant proteins were selected from class I in pink, class II in orange, and Rab family in grey).

**Figure S4. The distinctive case of GPAT4.** Left: Zoomed microscopy image of GPAT4 transfected HeLa WT in hypotonic media. White arrow points to LD with high targeting of GPAT4, other LDs have a lower protein intensity at their surface. Scale bar is 5 µm. Right: Individual droplet measurements of partition coefficient of GPAT4 in HeLa WT and SKO.

**Figure S5. Characterization of droplet incorporation in GERVs by FRAP experiments. (A)** TAG-in-media emulsion was complemented with Rhodamine-DPPE (at 1:7000 w:w Rho-DPPE:TAG). We performed FRAP experiments by photobleaching the whole droplet and measuring the recovery of Rho-PE: graph shows normalized intensity at the droplet surface according to time, with a zoomed graph in the inset showing partial recovery of the PLs signal. **(B)** TAG-NBD-in-media emulsion was used and the whole TAG droplet was photobleached. The normalized volume intensity in the droplet according to time shows a partial recovery of the NLs signal (followed by a decay due to photobleaching at long time). **(C)** DEGERVs were prepared with HNeu-GFP transfected cells. We performed FRAP on the ER membrane and on the whole aLD: a slower but consistent recovery was measured on LD as compared to ER, indicating that ER and LD can exchange protein.

**Figure S6. Confocal images of DEGERVs and comparison of partition coefficients in cells (WT and SKO) and DEGERVs. (A)** Confocal microscopy images for each protein with protein signal in green and ER lumen marker in blue. White arrows point to the GERV-embedded droplet. Scale bars are 5 µm. **(B)** Partition coefficients for our 17 proteins subset represented with mean value as a bar and standard deviation as gray whiskers (log scale), measured in HeLa WT (white bars), in HeLa SKO (colored bars), and in DEGERVs (striped bars). The average values of three independent experiments are represented as circle, square, and triangle gray symbols. Result of one-way ANOVA are shown on graph (see Table 2 for values), difference is non-significant when nothing is indicated.

**Figure S7. High-affinity proteins competition in DEGERVs. (A, B)** We analyzed the competition between high-affinity proteins: PLIN1/ACSL3, PLIN1/HPos, and PLIN1/HSD17B13. **(A)** Partition coefficients are normalized by the average partitioning of a protein transfected individually (left part of graph). We can visualize that, even though other proteins are not fully displaced, PLIN1 dominates the relocation to LDs when competing with ACSL3, HPos or HSD17B13. **(B, C)** For each couple of proteins, we show confocal images of an embedded droplet (left panel) and we compare the partition coefficient of both proteins when transfected individually (right panel; white bars, Fig. 2F), and upon co-transfection (right panel; colored bars) for HPos/PLIN1 (B), and PLIN1/HSD17B13 (C). Average values of three independent experiments are represented as circle, square and triangle gray symbols. All data points are represented in light gray (between 3 to 22 droplets for each experiment). Results of the unpaired t-test are shown on the graph (see Tables 3 and 4).

**Figure S8. DGAT2 is highly sensitive to crowding in HeLa cells. (A, C)** Confocal microscopy images of cells co-transfected with.HPos and DGAT2 (A) and PLIN1 and DGAT2 (C) were observed in normal media. Line profiles in green and red channels along the yellow line are shown below in (A). **(B, D)** Cells observed in hypotonic media. **(B)** Quantification of the partition coefficients of both proteins when transfected individually (white bars, Fig. 1E), and upon co-transfection (colored bars). Average values of three (resp. two for the co-transfection) independent experiments are represented as circle, square and triangle gray symbols. All data points are represented in light gray (between 73 to 80 droplets total, on 4 to 5 cells for each experiment). Result of unpaired t-test are shown on graph (see Tables 5 for detailed values). We observe a clear anticorrelation between P(HPos) and P(DGAT2) at low partition coefficients, revealing a competition for space at LD surface.

**Figure S9. Proteomics and immunofluorescence images of preadipocytes and adipocytes. (A, B)** Different volcano plot representation of proteomics data: relative protein intensity in LD fraction in preadipocytes *versus* adipocytes (10 days) (A) and relative protein intensity in the PNS fraction in preadipocytes *versus* adipocytes (10 days) (B). Statistical significance -log_10_(P-Value) is plotted as a function of the fold change, with class I proteins in pink, class II in orange and Rab family proteins in blue. Proteins above the dashed line correspond to a P-Value < 0.05. **(C, D)** Immunofluorescence images of endogenously expressed DGAT2 (green) in preadipocytes without oleate (C) and in adipocytes exposed to 1 mM Na^+^oleate for 5h (D). **(E-H)** Immunofluorescence images of endogenously expressed ACSL3 (green) in preadipocytes without Na^+^oleate (E), adipocytes without oleate (F), in preadipocytes exposed to 500 µM oleate for 8h (G) and in adipocytes exposed to 1 mM Na^+^oleate for 5h (H). LDs are labelled with LipidToxRed (blue).
