## Supplementary figures and images for "Steric Repulsion Counteracts ER-to-Lipid Droplet Protein Movement"

### Supplementary figures

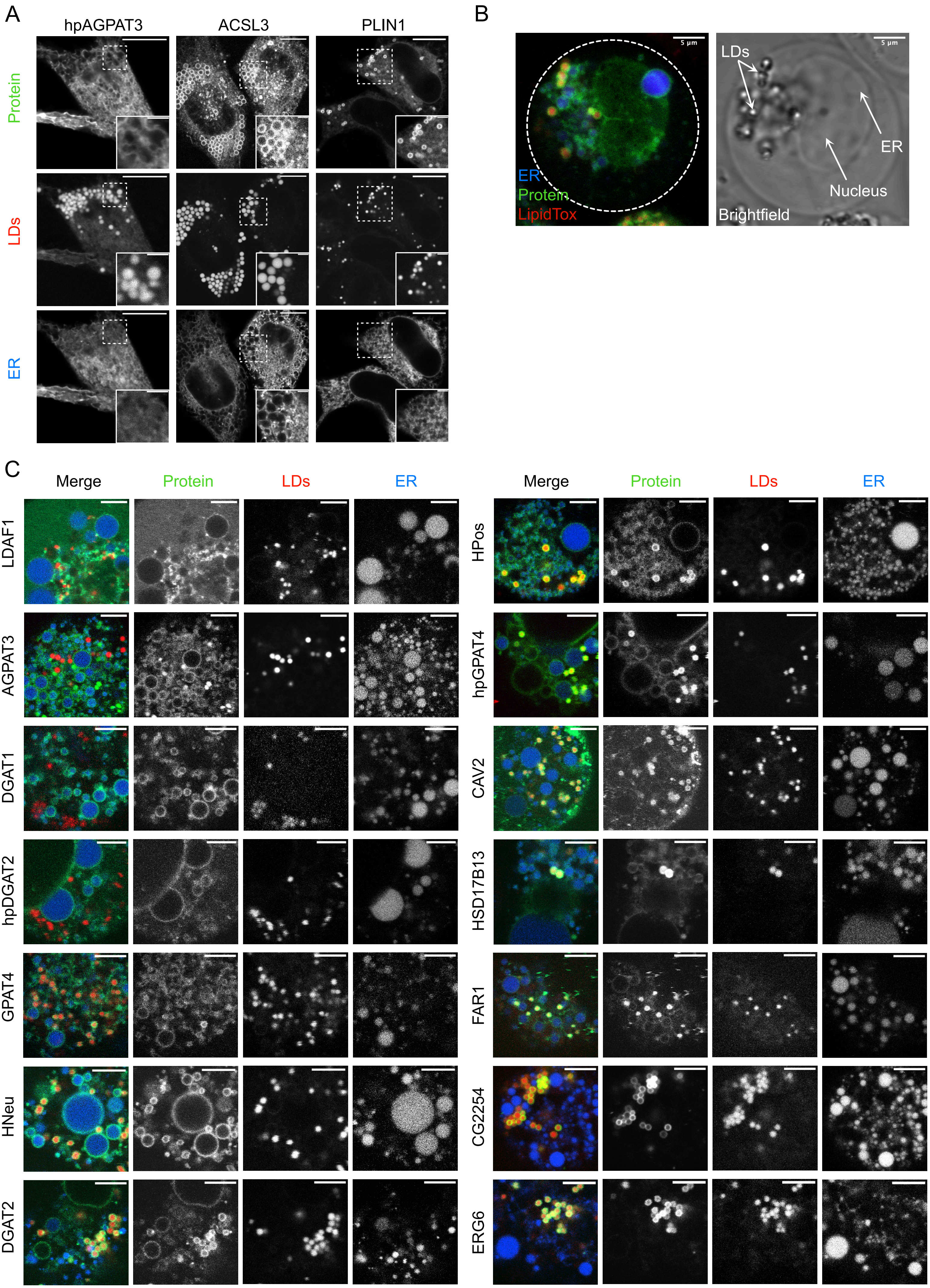

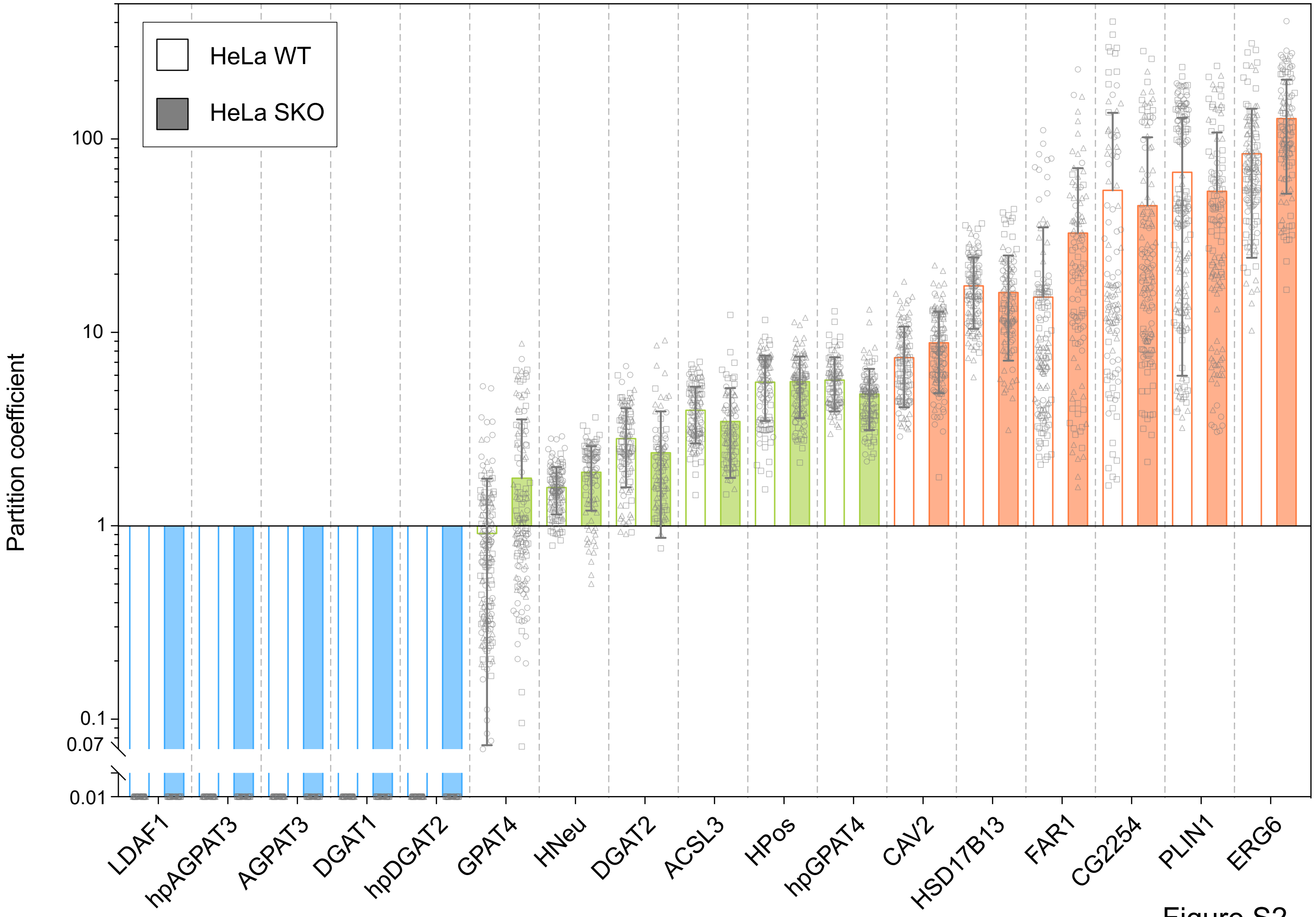

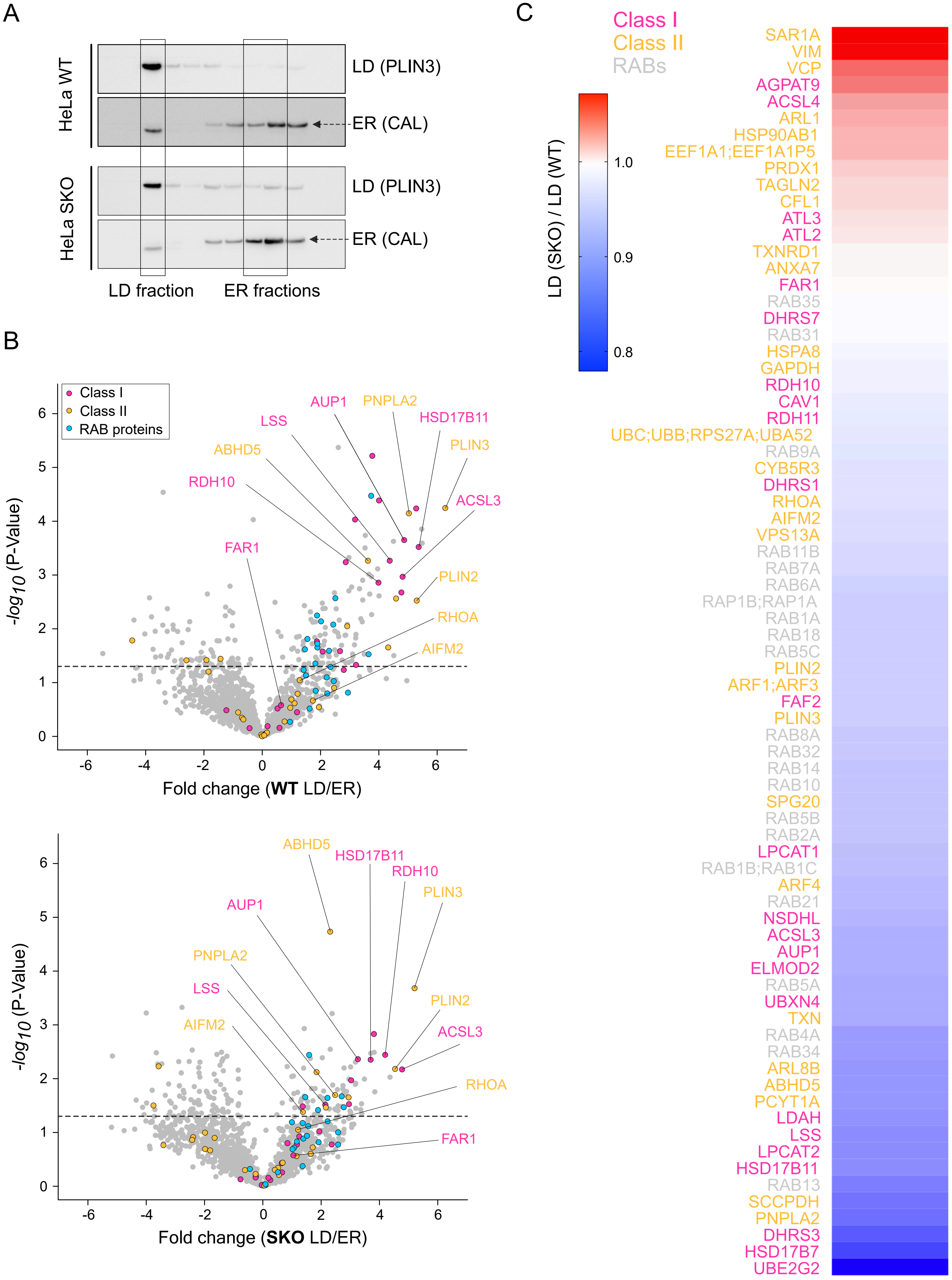

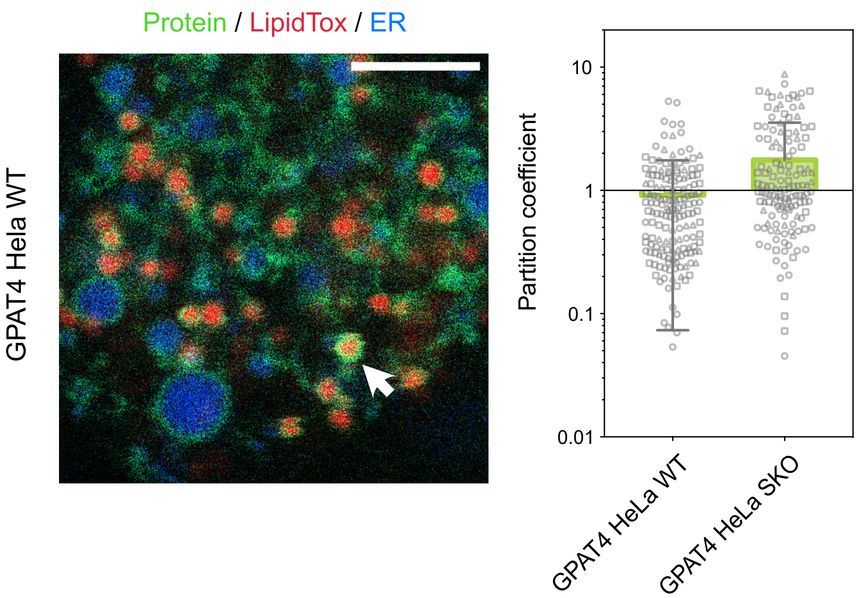

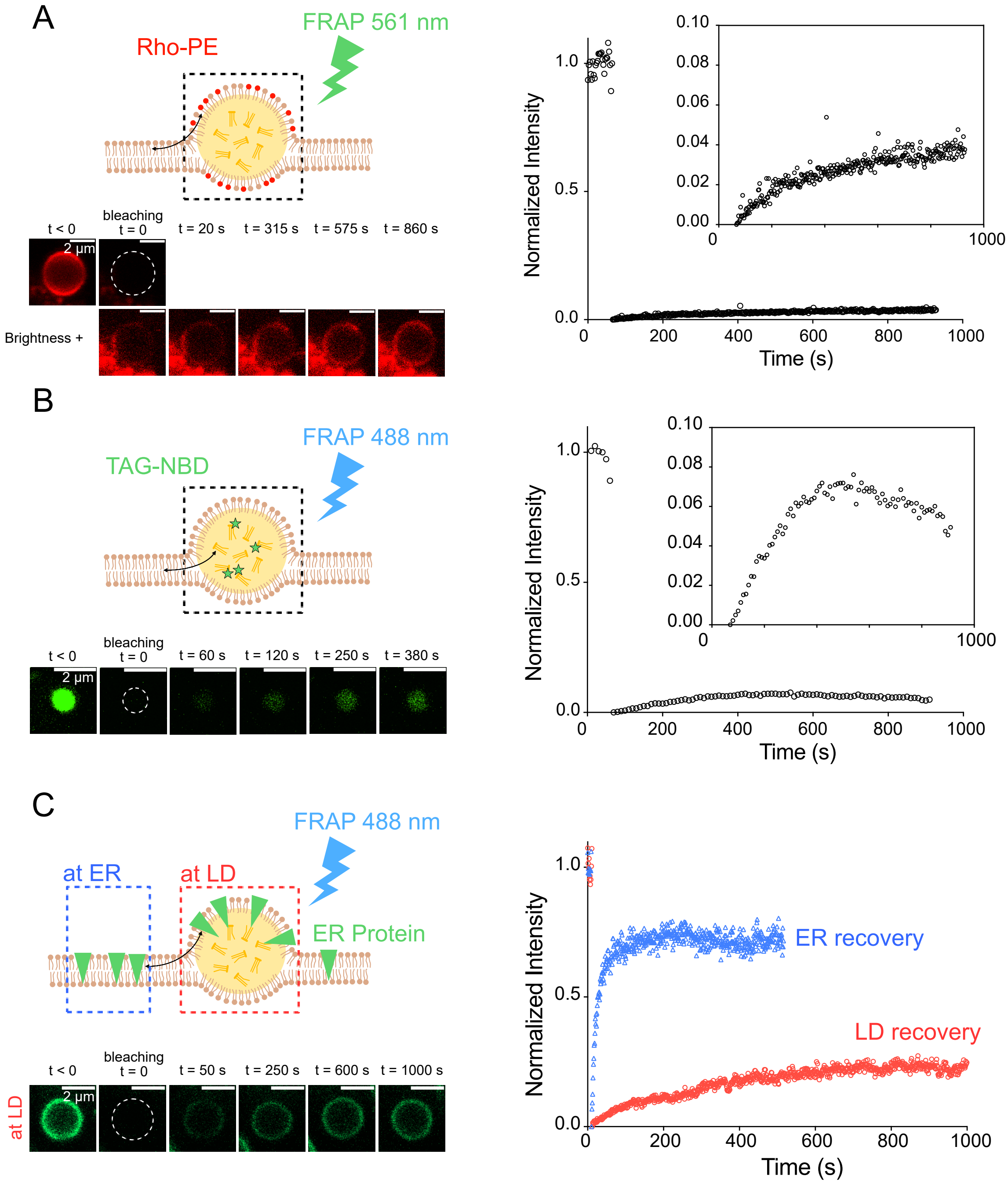

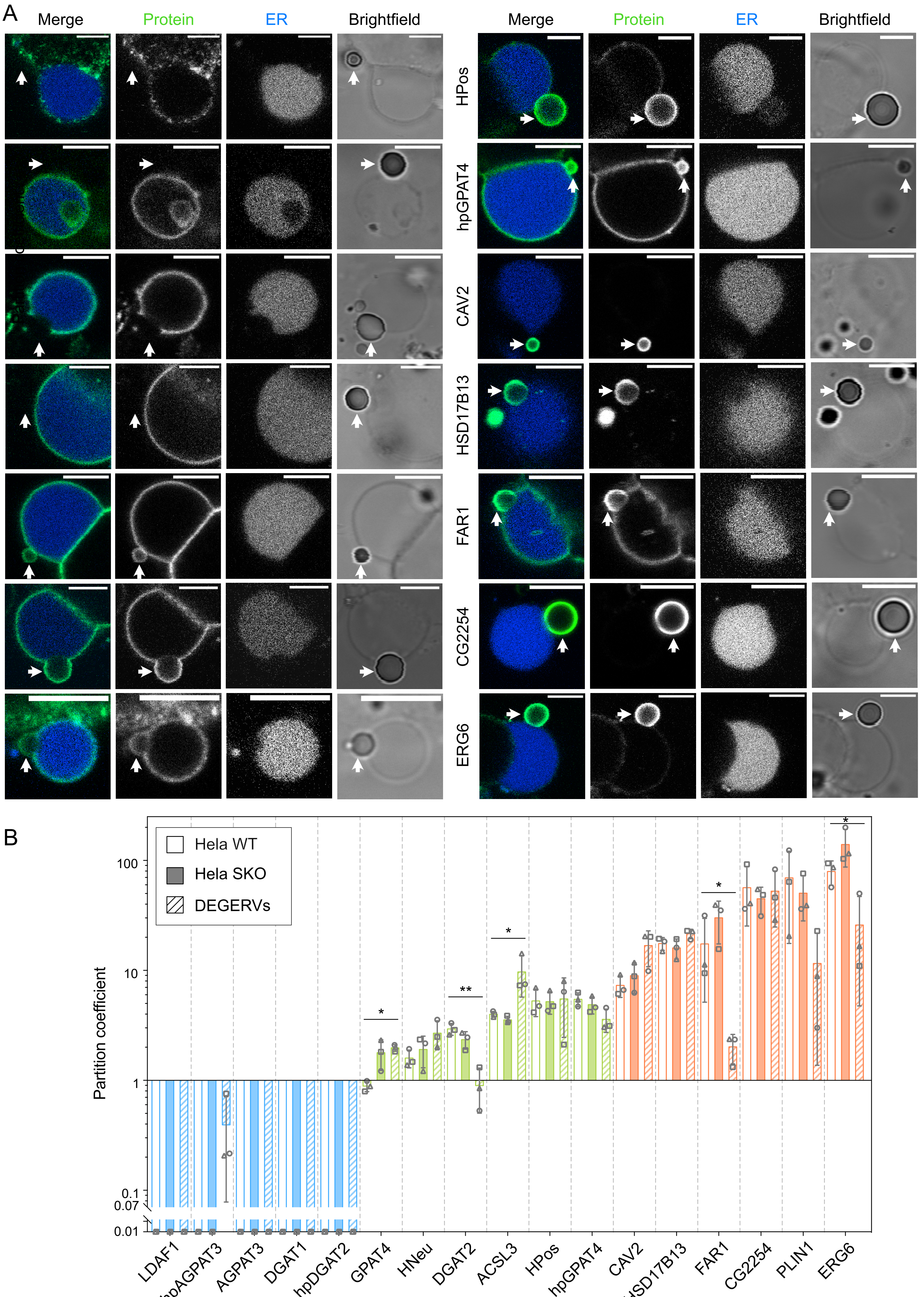

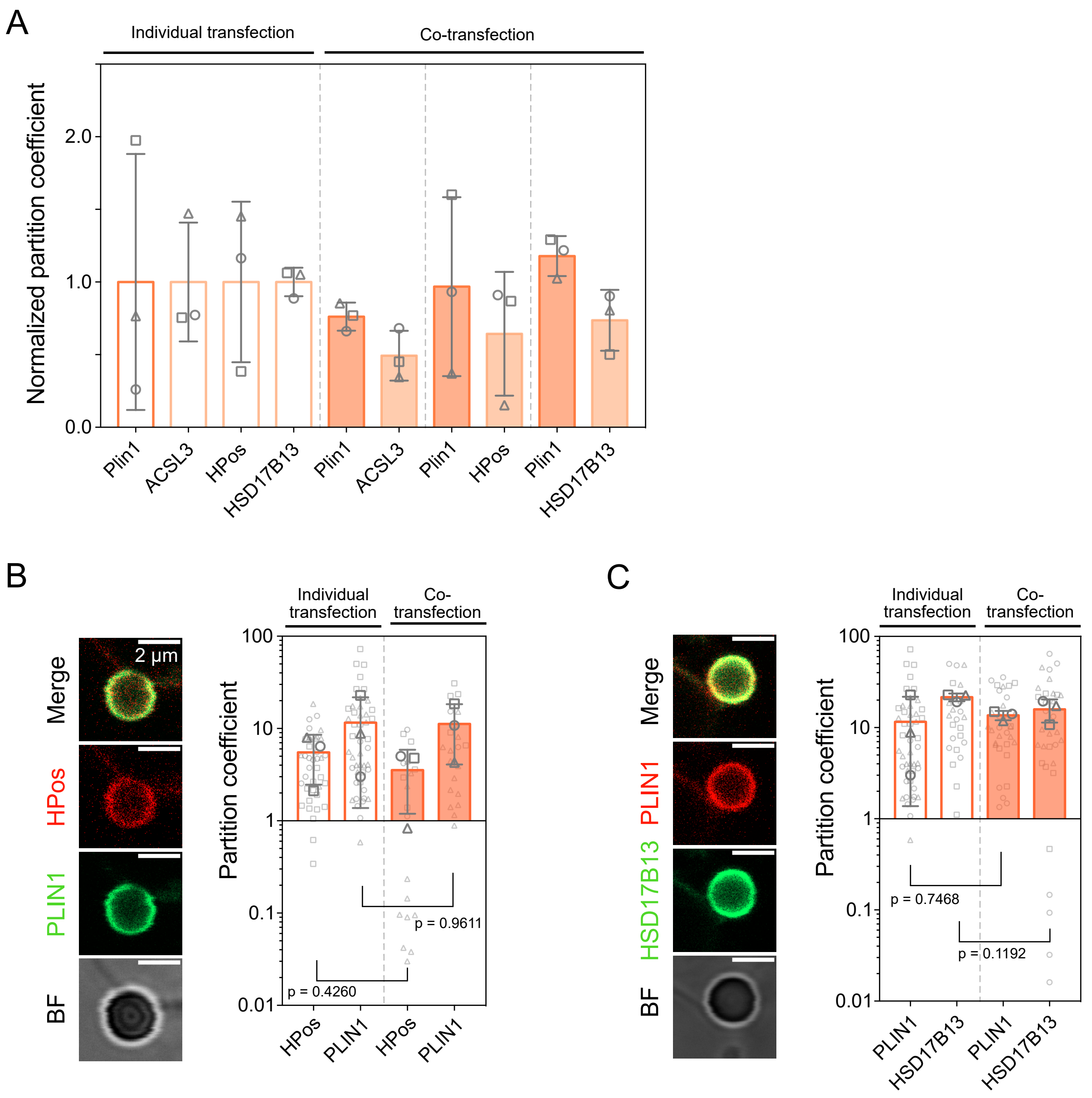

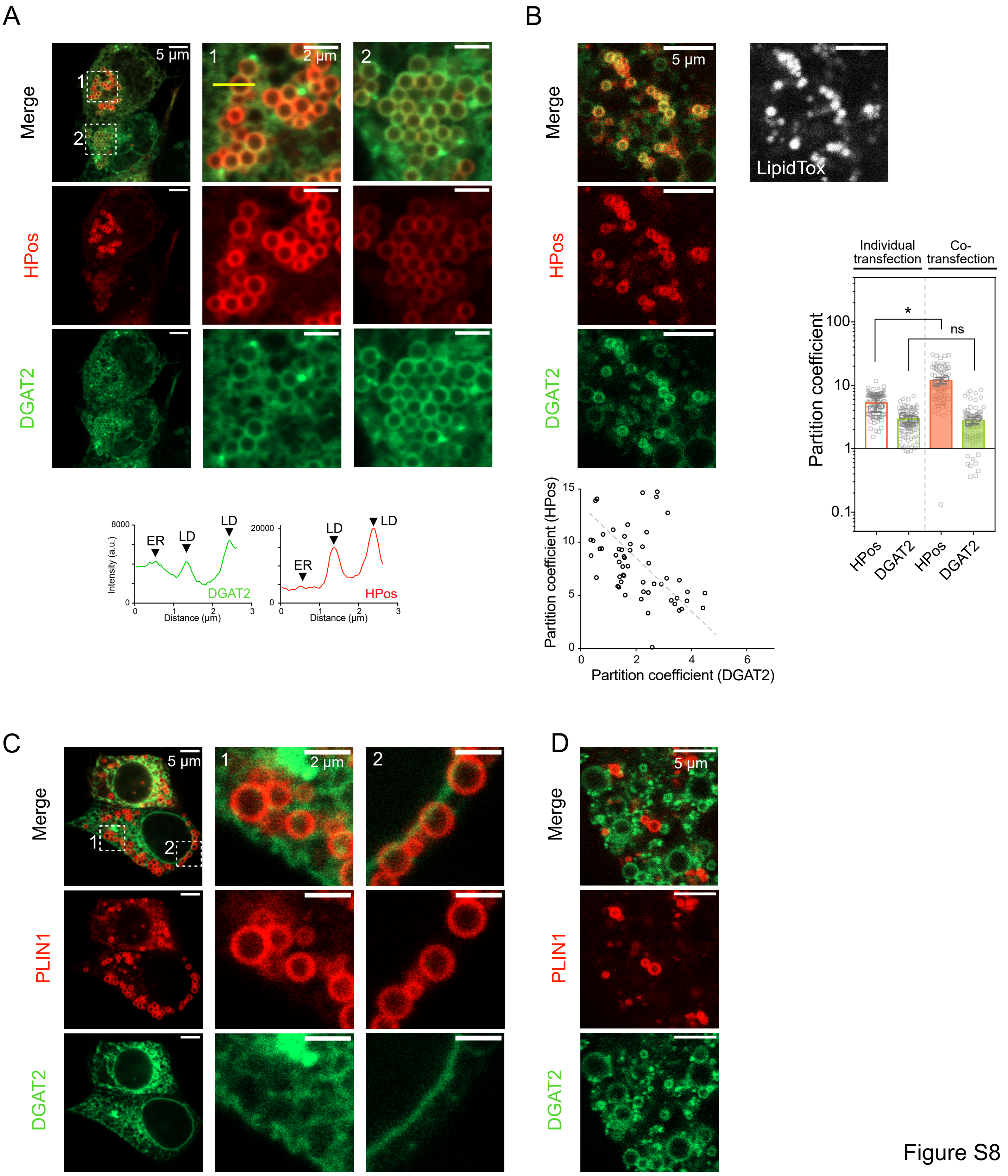

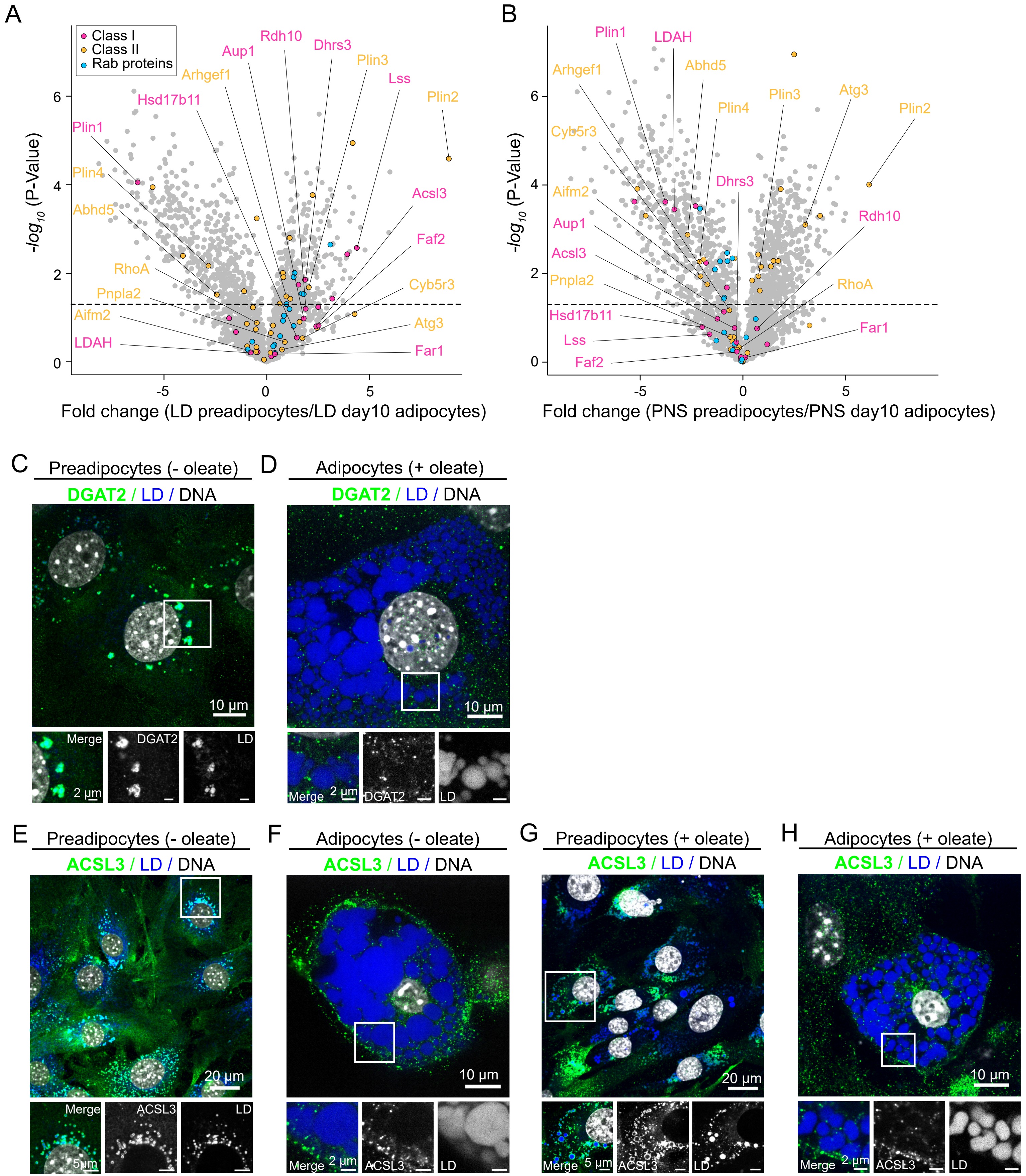
